# Analysis of Infiltrating Immune Cells Following Intervertebral Disc Injury Reveals Recruitment of Gamma-Delta (γδ) T cells in Female Mice

**DOI:** 10.1101/2024.03.01.582950

**Authors:** Sade W. Clayton, Remy E. Walk, Laura Mpofu, Garrett W.D. Easson, Simon Y. Tang

## Abstract

Inadequate repair of injured intervertebral discs (IVD) leads to degeneration and contributes to low back pain. Infiltrating immune cells into damaged musculoskeletal tissues are critical mediators of repair, yet little is known about their identities, roles, and temporal regulation following IVD injury. By analyzing longitudinal changes in gene expression, tissue morphology, and the dynamics of infiltrating immune cells following injury, we characterize sex-specific differences in immune cell populations and identify the involvement of previously unreported immune cell types, γδ and NKT cells. Cd3+Cd4-Cd8-T cells are the largest infiltrating lymphocyte population with injury, and we identified the presence of γδ T cells in this population in female mice specifically, and NKT cells in males. Injury-mediated IVD degeneration was prevalent in both sexes, but more severe in males. Sex-specific degeneration may be associated with the differential immune response since γδ T cells have potent anti-inflammatory roles and may mediate IVD repair.

Graphical Abstract:
Schematic of the workflow to obtain longitudinal analyses of the acute IVD injury response Injured caudal IVDs, CC5/6-CC9/10, were bilaterally punctured with a 30G needle to induce a traumatic injury. Injured IVDs and neighboring uninjured internal Control IVDs, CC12/13-CC16/17, were isolated during the acute injury response every 2-3 days until 21 days post injury (dpi) and a chronic injury time point at 42 dpi in female mice. Male mice samples were only collected at key injury response time points (red numbers): 3, 7, 12, 19, and 42 dpi. Longitudinal analyses of the temporal regulation of immune cell gene expression and cell infiltration were measured with qPCR, flow cytometry, immune fluorescence, and histology analyses to identify a sex-divergent immune response.

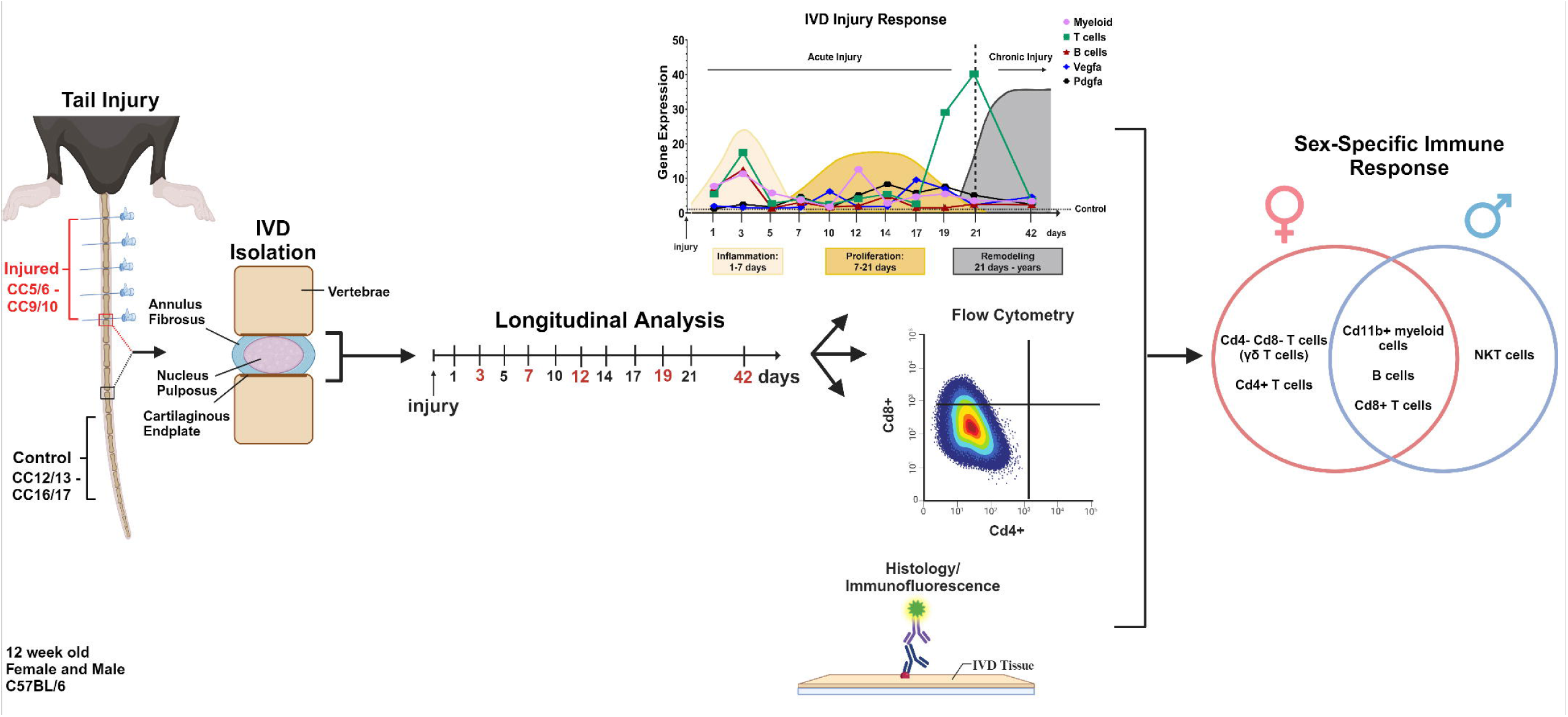

## Introduction

Intervertebral discs (IVD) are spinal, fibrocartilaginous joints that are prone to cumulative injuries overtime which ultimately lead to compounding, progressive degeneration^1,2^. IVD degeneration is a major contributor to low back pain and damaged IVDs repair poorly^1,3,4^. The biological repair of injured tissue is a dynamic, multifactorial process that is essential for tissue health and dysregulation leads to chronic disorders^5^. The IVD consists of three tissue types, the fibrocartilaginous annulus fibrosus, gelatinous nucleus pulpous, and the cartilaginous endplate, that exist in an aneural, avascular, hypoxic environment^6^. The etiology of why IVD injuries lead to degeneration instead of tissue repair is poorly understood, but studies have suggested the heterogeneous structure of the IVD as a contributor^6^. Upon injury, the IVD becomes chronically vascularized, innervated, and remains in a proinflammatory state, which is thought to be the major generator of IVD-related back pain^7,8^. Many studies have investigated the chronic effects of IVD injury, but few have investigated the immediate changes that occur acutely post injury. The acute injury response (up to 2 to 4 weeks post injury) is a critical, tissue-dependent time window where essential growth factors and immune cells drive the tissue repair process^9–11^. Perturbations in the acute injury response leads to chronic disrepair, but the association between the IVD acute injury response and how this process contributes to chronic degeneration is understudied. There is a critical need to understand the cell types and signaling pathways that define the acute injury response to identify targetable factors to increase the regenerative capacity of the IVD.

The regenerative capacity of musculoskeletal tissues is attributed to the efficacy of injured tissue to undergo complete biological repair after injury. Tissue repair is a regulated temporal process characterized by the progression of three sequential, semi-overlapping stages: Inflammation, Proliferation, and Remodeling^10,12–14^. Inflammation involves rapidly infiltrating immune cells into the injury site to survey for pathogens, clear cell debris, and secrete cytokines to stimulate the next stage, Proliferation. The Proliferation stage replenishes cells that were lost by bringing nutrients to the injury site through stimulating angiogenesis via Vegf and Pdgf pathways as inflammation wanes. Lastly, the Remodeling stage stimulates extracellular matrix remodeling of replenished cells to restore tissue mechanical strength. Infiltrating immune cells have been shown to be critical mediators of not only the inflammation stage but also essential throughout the acute stages of the injury response in musculoskeletal tissues and the skin^15–18^. Both myeloid cell types, such as macrophages, monocytes and neutrophils, and lymphocyte cell types, such as B and T cells, rapidly infiltrate both regenerative musculoskeletal tissues such as bone, muscle, and those only capable of limited functional repair like cartilage and tendon during acute injury time points. Moreover, the immune system has a critical role in the repair response since inhibition of these immune cells lead to delayed or completely disrupted tissue repair^12,15,19,20^. The importance of identifying and understanding the immune cell response during acute injury is clear, yet the IVD is one of the few remaining musculoskeletal tissues where crosstalk between the immune system the acute injury response has yet to be characterized.

In this study, we employ a combinatory approach by using qPCR, flow cytometry, histology, and immunofluorescence to identify the longitudinal changes in the identities, gene expression, and localization of immune cells during the acute IVD injury response in 12-week old C57BL/6 mice. We hypothesize that the acute infiltration of immune cells into injured IVDs is a tightly controlled temporal process that is associated with degeneration..

## Methods

### Animals

12-week old female (n= 43) and male (n= 21) C57BL/6J (#000664) mice were purchased from the Jackson Laboratories and housed in a mouse facility in ventilated cages under standard laboratory conditions where chow diet and water were available at libitum. Procedures were performed within seven days of animal deliveries to ensure animals were aged-matched. qPCR, histology, and immunofluorescence experiments were conducted using the same animals. Flow cytometry was performed by using separated animals per experiment. Institutional Animal Care and Use Committee protocols were established and approved before animal usage and conformed to the National Institutes of Health Guide for the care and use of laboratory animals.

### Intervertebral Disc Injury

Five Control (non-injured) and five Injured (bilateral, 30G needle puncture) tail IVDs were extracted for longitudinal analysis (qPCR) of the acute injury response every 2-3 days post injury until 21 days post injury (dpi) and a chronic injury time point of 42 dpi in 12-week old female C57BL/6J mice. A similar procedure was conducted in 12-week old male BL6 mice, but only at select, key time points: 3,7,12, 19 and 42 dpi (**Figure 1**). Needle punctures were performed by using digital palpation to identify coccygeal (CC) regions CC5/6 through CC9/10 in mice tails, marking the levels with a Sharpie, puncturing each level with a sterile 30G needle, and confirming the accuracy of each puncture with X-ray images (Faxitron UltraFocus 100, Hologic). Mice were anesthetized with vaporized 3% isoflurane/oxygen during the procedure and given a onetime subcutaneous injection of 1mg/mL carprofen at 5mg/kg/mouse as analgesia immediately after needle punctures. Mice were returned to the mouse facility and monitored every 24 hours until the correct disc extraction time points. For tail disc extractions, mice were euthanized at the aforementioned time points in a CO2 chamber with 3% CO2 for 5 minutes and a 2 minute dwell time. Sacrificed mice were submerged in 70% ethanol for 2 minutes and the sacral and caudal spine removed. All extraneous tissues were removed from the tail until only the vertebrae and discs remained, and CC5/6 through CC9/10 and CC12/13 through CC16/17 were isolated for the Injured and Control groups, respectively. For each treatment group, three discs were randomly pooled together in an Eppendorf tube, snap frozen in liquid nitrogen and kept at -80 degrees Celsius for future RNA extraction. The remaining two discs were used for histology.

**Figure 1:**
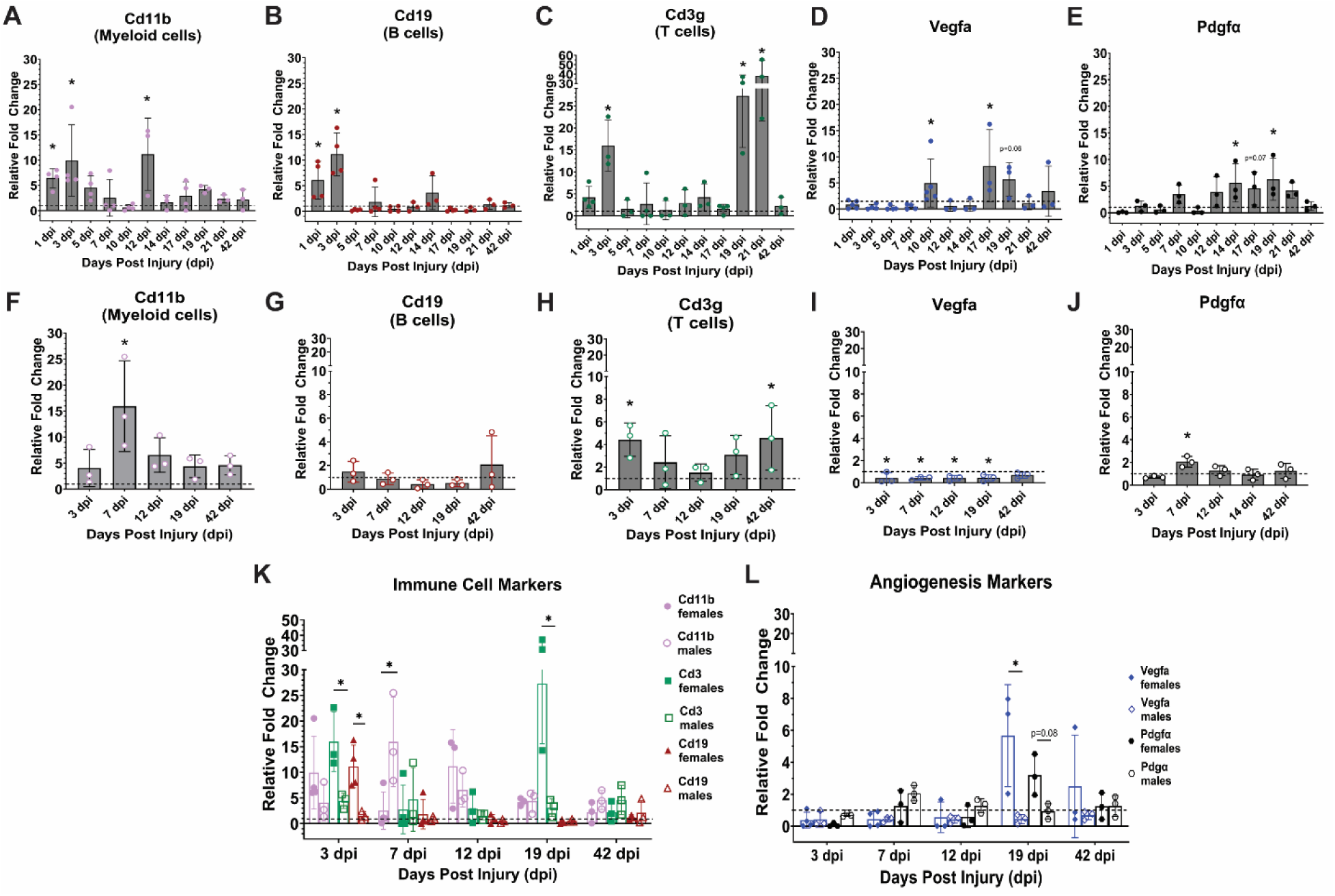
Gene marker regulation of immune cells is temporally divergent between female and male mice during the acute IVD injury response. Gene markers for immune cells that broadly identify subtypes of (A) myeloid cells (*Cd11b*), (B) B cells (*Cd19*), and (C) T cells (*Cd3g*) were used for qPCR analysis of the temporal regulation these cell types over the course of the acute IVD injury response in Injured IVDs (bars) in comparison to Controls (dotted black line) in females. (D) *Pdgfa* and (E) *Vegfa* gene expression was measured longitudinally to denote the wanning of the Inflammation stage and the time course of the Proliferation stage during the female acute IVD injury response. Gene regulation of (F) *Cd11b*, (G) *Cd19*, and (H) *Cd3g*, (I) *Pdgfa*, and (J) *Vegfa* was measured in males at the key time points of dynamic gene regulation based on the female longitudinal analyses: 3, 7, 12, 19, 42 dpi. A comparative analysis of the sex-specific differences in gene regulation of (K) the immune cells gene markers and (L) angiogenic gene markers at the key time points show stark differences in the temporal regulation of gene markers between sexes. A-J: asterisks denote statistical significance between control and injured samples. K and L: asterisks denote statistical significance between sexes. Exact p values are listed in Table 2 and Table 3.

### RNA Isolation and Reverse Transcriptase Quantitative Polymerase Chain Reaction (qPCR)

Pooled, frozen discs were homogenized by being placed in a Mikro-Dismembrator U (Sartorious Group) that grinded the tissues at 2000 rpm for 40 seconds. Cells were lysed by using TRIzol reagent (Fisher Scientific) and total RNA extracted using a Direct-zol RNA MicroPrep Kit (Zymo Research). RNA concentrations were measured with a spectrophotometer (Nano Drop) and 2 ng of RNA per sample was used in each qPCR reaction according to instruction in the ZymoScript Onestep RT-qPCR Kit (Zymo Research). qPCR was conducted using a QuantStudio3 (Applied Biosystems) to measure crossing point values, which were converted to relative fold changes by using ΔΔCt method. Gene markers for infiltrating myeloid cells (*Cd11b*), B cells (*Cd19*), T cells (*Cd3g*) and angiogenic factors *Vegfa* and *Pdgf*α were analyzed and *HPRT/GAPDH* were used as a normalization controls, n= 3-5. “n” = biological replicates (mice). Primer sequences for qPCR are shown in **Table S1**.

### Histology

Two discs from randomized levels from the Control and Injured groups were used for Safranin O and Fast Green staining (Fisher Scientific) to identify changes in IVD tissue morphology. Tissues were decalcified for 3-4 weeks in 14% EDTA at 4 degrees Celsius. The tissues were then placed in 30% sucrose overnight at 4 degrees Celsius and then changed to graded 30% sucrose: OCT solutions for 1 hour until reaching 100% OCT. Levels were embedded in OCT and 10μm sections were collected using a cryostat (Lecia). Slides containing sagittal, midline sections from 3,7,12, 19 and 42 dpi were fixed with 4% PFA for 10 minutes before performing histological staining. Slides were stained with 0.1% Safranin O to identify glycosaminoglycans and 0.05% Fast Green as a counter stain and then imaged with an Olympus BX51 microscope using a 4X objective.

### Flow Cytometry

Approximately 23,000 cells from Controls and 18,000 cells from Injured IVDs were collected for flow cytometry by pooling 5 IVDs for each treatment group from each mouse in a total of three mice to collect 15 IVDs/ flow cytometry run. Pooled IVDs were isolated and rendered into a single cell suspension by using four serial digestions of 2mg/mL Collagenase Type II (Gibco) in DMEM at 37 degrees Celsius for 30 minutes each. After resuspension in FACS buffer (0.5% BSA, 2mM sodium azide, 2mM EDTA in 1X PBS), cells were counted using a hemacytometer and then treated with an Fc blocking buffer containing anti-Cd16/32 (BD Biosciences, 1:500) for 20 minutes at 4 degrees Celsius before a 30 minute incubation with antibody master mixes for the myeloid or lymphoid panels at 4 degrees Celsius. The myeloid panel consisted of Cd45-APCCy7 (BD Biosciences), Cd11b-BV605 (Biolegend), Ly6G-BV510 (Biolegend), Ly6C-PE (Biolegend), F4/80-PE Cy7 (Biolegend), Cd11c-Fitc (Biolegend), Cd206-Alexa 700 (Invitrogen), Cd80-PE Cy5 (Invitrogen), Siglec F-BV421 (BD Biosciences), and 7AAD (Thermofisher) as a live dead stain. The lymphoid panel consisted of Cd45-FITC (eBioscience), Cd3-BV510 (Biolegend), Cd4-PE (eBioscience), Cd8-APC (eBioscience), Cd19-PECy7 (eBioscience), Nk1.1-BV605 (eBioscience) and 7AAD. For compensation, the spleen from one of the mice was isolated, homogenized into a single cell suspension, treated with Red Blood Cell Lysing Buffer Hybri-Max™ (Sigma Aldrich) and aliquoted to 1 million cells/ tube in FACS buffer to be treated with individual antibodies from each panel to serve as single stained controls. After staining, cells were identified by using a LSRFortessa (BD Biosciences) analyzer and FloJo 10.0 software for cell population gating.

### Immunofluorescence

Midline, sagittal sections from 3, 7, 12, 19 and 42 dpi samples from both sexes were fixed with 4% PFA for 10 minutes and then permeabilized with 0.3% Triton X in TBS for 10 minutes. Sections were washed and then blocked using 1% Goat serum in TBS for one hour at room temperature before adding primary antibodies to detect Cd45 (Cell Signaling, 1:500) and Cd11b (Invitrogen, 1:350) or Cd3 (Invitrogen, 1:100) and TCRγδ (eBiosience, 1:80) either overnight at 4 degrees Celsius or for 4 hours at room temperature. An anti-mouse Armenian hamster Alexa 555, rat Alexa 488, rabbit Alexa 488, and rat Alexa 594 secondary was added at a dilution of 1:250 for one hour at room temperature dependent upon the primary antibodies. Slides were treated with Hoechst 33258 (Invitrogen) at a dilution of 1:1000 before being mounted and imaged with a Leica Di8 laser scanning confocal microscope with a 10x objective. 2 to 3 sequential sections per sample were imaged, quantified, and averaged for statistical analyses.

### Statistical Analyses

All statistical analyses were performed using GraphPad Prism version 10.2.0. The assumptions for parametric tests were checked to verify none were violated before statistical tests were ran and results interpreted. For female and male longitudinal gene expression data, differences between the two factors, injury (paired) and dpi (independent), and the interaction term were analyzed by a mixed design two-way ANOVA followed by Holm-Šídák’s multiple comparisons test. The post hoc analyses are detailed in **Table S2** and **Table S3**. For comparative analyses between female and male gene expression, differences between the two independent factors, sex and gene, and the interaction term at each key time point (3,7,12,19, and 42 dpi) were analyzed by two-way ANOVA followed by Holm-Šídák’s multiple comparisons (**Table S4**). The difference in the proportions of infiltrating cells between females and males observed with flow cytometry was measured with chi-squared tests. For immunofluorescence (Cd45 and Cd11b) and histology, a mixed design two-way ANOVA followed by Fisher’s LSD multiple comparisons test was used to measure differences between the two independent factors, sex (independent) and injury (paired), and determine the interaction term. For Cd3 and TCRγδ, differences were measured using unpaired Student’s T tests. A p value < 0.05 was considered statistically significant and an asterisk denotes significance. “ns” denotes not statistically significant. Degeneration scoring was performed blinded by three to four people from the laboratory for each sample and the average score for the grading was used.

## Results

### Gene marker regulation of immune cells is temporally divergent between female and male mice during the acute IVD injury response

To determine the longitudinal regulation of immune cell genes during acute injury time points, we used primers to measure changes of gene expression in Control (dashed line) and Injured (bars) IVDs from 1 to 42 dpi for myeloid cells (**Figure 1A**), B cells (**Figure 1B**), and T cells (**Figure 1C**) with qPCR in female mice. Myeloid cell gene expression, marked by Cd11b, oscillates after injury, with upregulation at 1 dpi, 3 dpi, and then is up-regulated again at 12 dpi. B cells, marked by *Cd19,* is only upregulated at 3 dpi. *Cd3g* shows a distinct oscillatory pattern for T cells where expression is upregulated early in the injury response at 3 dpi and a robust increase in gene regulation returns much later in the acute injury time points than any other immune cells at 19 and 21 dpi.

Angiogenesis is a critical component of the repair response during the wanning of the Inflammation phase and beginning of the Proliferation phase in injured tissues ^21^; therefore, we measured longitudinal gene expression changes for *Vegfa* (**Figure 1D**), and *Pdgf*α (**Figure 1E**). The Vegfa and Pdgfα signaling pathways are potent regulators of angiogenesis during repair of musculoskeletal tissues^22,23^. *Pdgf*α has a narrow window of upregulation from 14 to 19 dpi while *Vegfa* increase in expression at 10 dpi but is not upregulated again until 17 to 19 dpi.

The longitudinal expression patterns of the immune and angiogenic gene regulation revealed 5 inflection points of gene expression during IVD injury response in females: 3 dpi as the peak time of inflammation, 7 dpi as the wanning of inflammation and the beginning of angiogenesis regulation, 12 dpi as the return of myeloid cell marker expression, 19 dpi as the return of T cell marker expression, and 42 dpi as a chronic injury time point where both immune and angiogenic gene regulation has resolved (**Figures 1A-2E**). The chronic injury response is also a tissue-dependent time window but has been defined in other musculoskeletal tissues as beginning 4 to 6 weeks post injury^11^. At these key time points, we measured myeloid cells (**Figure 1F**), B cells (**Figure 1G**), T cells (**Figure 1H**), *Vegfa* (**Figure 1I**), and *Pdgf*α (**Figure 1J**) in 12-week old male mice. Males have a less dynamic and distinct temporal regulation of these gene markers than females where their acute injury response is characterized by *Cd11b* (**Figure 1F**) and *Pdgf*α (**Figure 1J**) regulation and an increase in Cd3g expression during chronic injury at 42 dpi (**Figure 1H**). Comparative analyses of immune cell genes (**Figure 1K**) and angiogenesis genes between sexes (**Figure 1L**) revealed the downregulation of myeloid, B, and T cells gene markers at 3 dpi, an upregulation of *Cd11b* at 7 dpi, and a stark down regulation of *Cd3* at 19 dpi in males when compared to injured female IVDs. The regulation of angiogenic factors is downregulated in males at 19 dpi as well.

### Males IVDs exhibit more severe degeneration following injury

Injured IVDs progressively degenerate over time and rarely undergo repair to complete functional restoration^24^. To determine the sex-specific differences in the time of progression and magnitude of degenerative changes over the key longitudinal time points, we stained midsagittal frozen sections of Control and Injured IVDs from 3 dpi (**Figure 2A**), 7 dpi (**Figure 2C**), 12 dpi (**Figure 2E**), 19 dpi (**Figure 2G**) and 42 dpi (**Figure 2I**) with Safranin O to visualize glycosaminoglycan content and tissue morphology. Degeneration scoring was performed by adhering to the standardized histopathology scoring system^25^. IVDs at 3 dpi have moderate degeneration with injury independent of sex (**Figure 2A, B**); however injured males exhibit accelerated degeneration compared to females between 7 dpi and 12 dpi (**Figure 2D, F**), though there is no sex-related difference in degeneration at 19 dpi (**Figure 2H**). We then analyzed samples at 42 dpi, a chronic injury time point that falls into the remodeling stage, to assess if either sex has the capability to undergo tissue repair, which we define as injured IVDs showing no signs of the needle puncture injury and resembling uninjured controls. At 42 dpi, female IVDs have similar degeneration scores as 19 dpi, while male mice have elevated degenerative scores that are in the moderate range (**Figure 2I, J**). There is no repair in either sex, but males have higher degeneration scores at 42 dpi.

**Figure 2:**
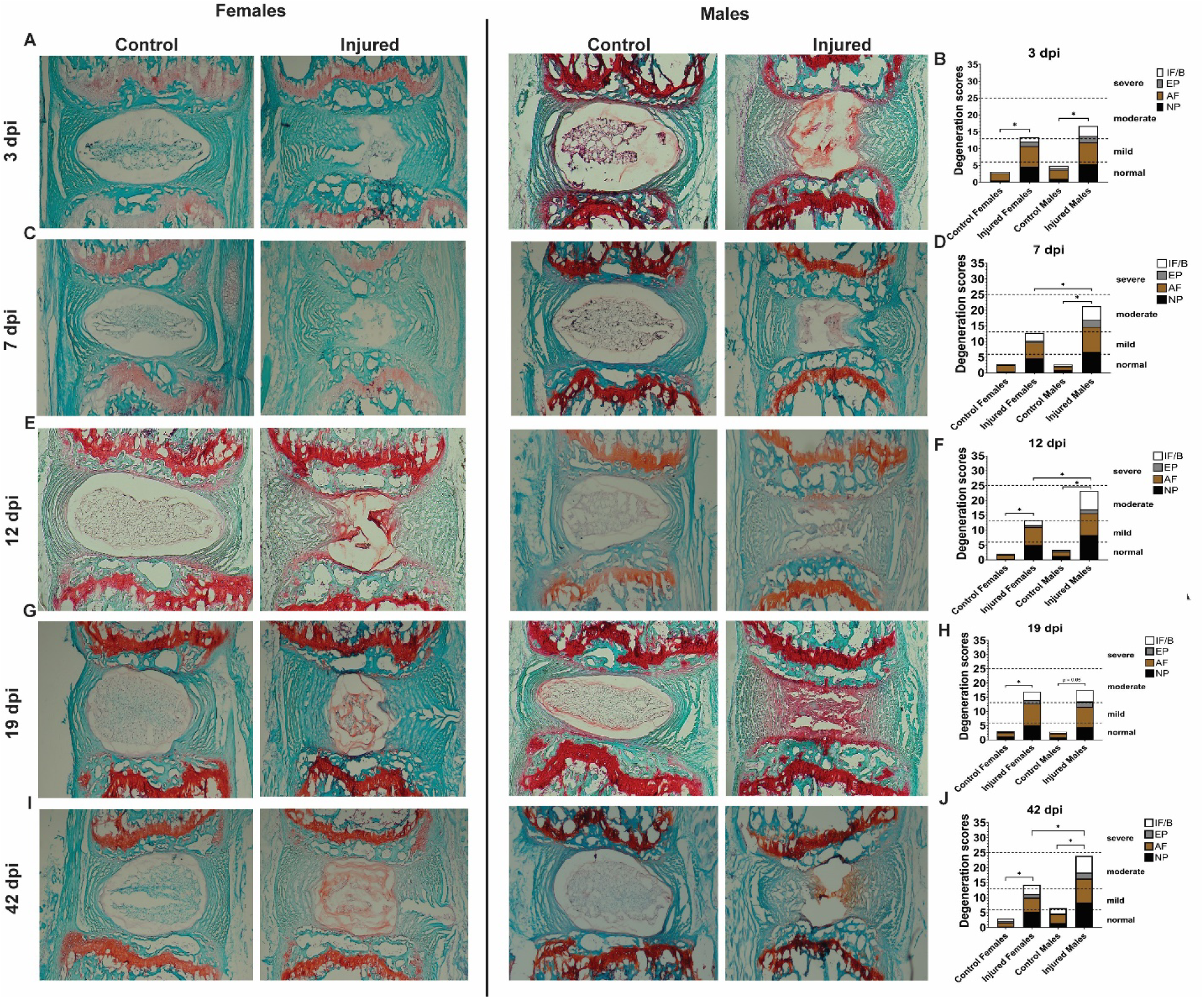
Males IVDs exhibit more severe degeneration following injury. Midsagittal sections of Control and Injured IVDs from both sexes at (A) 3 dpi, (C) 7 dpi, (E) 12 dpi, (G) 19 dpi, and (I) 42 dpi were stained with Safranin O and Fast Green dyes to visualize IVD tissue morphology. Degenerative scores were measured and quantified for (B) 3 dpi, (D) 7 dpi, (F) 12 dpi, (H) 19 dpi, and (J) 42 dpi.

### Males have higher percentages of responding Cd11b+ myeloid cells with injury at 3 dpi

We wanted to identify the association between the observed temporal differences in the immune cell gene regulation and tissue morphology with the presence of immune cell sub-types in injured IVD tissue. 3 dpi was the timepoint with the most abundant differential expression of immune cells genes between females and males; therefore, we utilized flow cytometry to determine the differences in the innate immunity response by profiling Cd11b+ antigen presenting cells (APC) and phagocytes indicated to mediate acute inflammation and repair^26^. There is an increase in Cd45+ immune cells and Cd11b+ myeloid cells in response to injury in both sexes. Females have a higher percentage of resident Cd11b+ cells and males have a larger increase in Cd11b+ cells with injury (**Figure S1**). The greatest proportion of resident myeloid cells in females are Ly6c+ monocytes (**Figure 3A**), while Ly6g+ neutrophils are the majority in males (**Figure 3C**). With injury, monocytes dominate the myeloid cell response in both sexes and there is a shift in the percentage of responding neutrophils dependent upon sex (**Figure 3B’, D’**).

**Figure 3:**
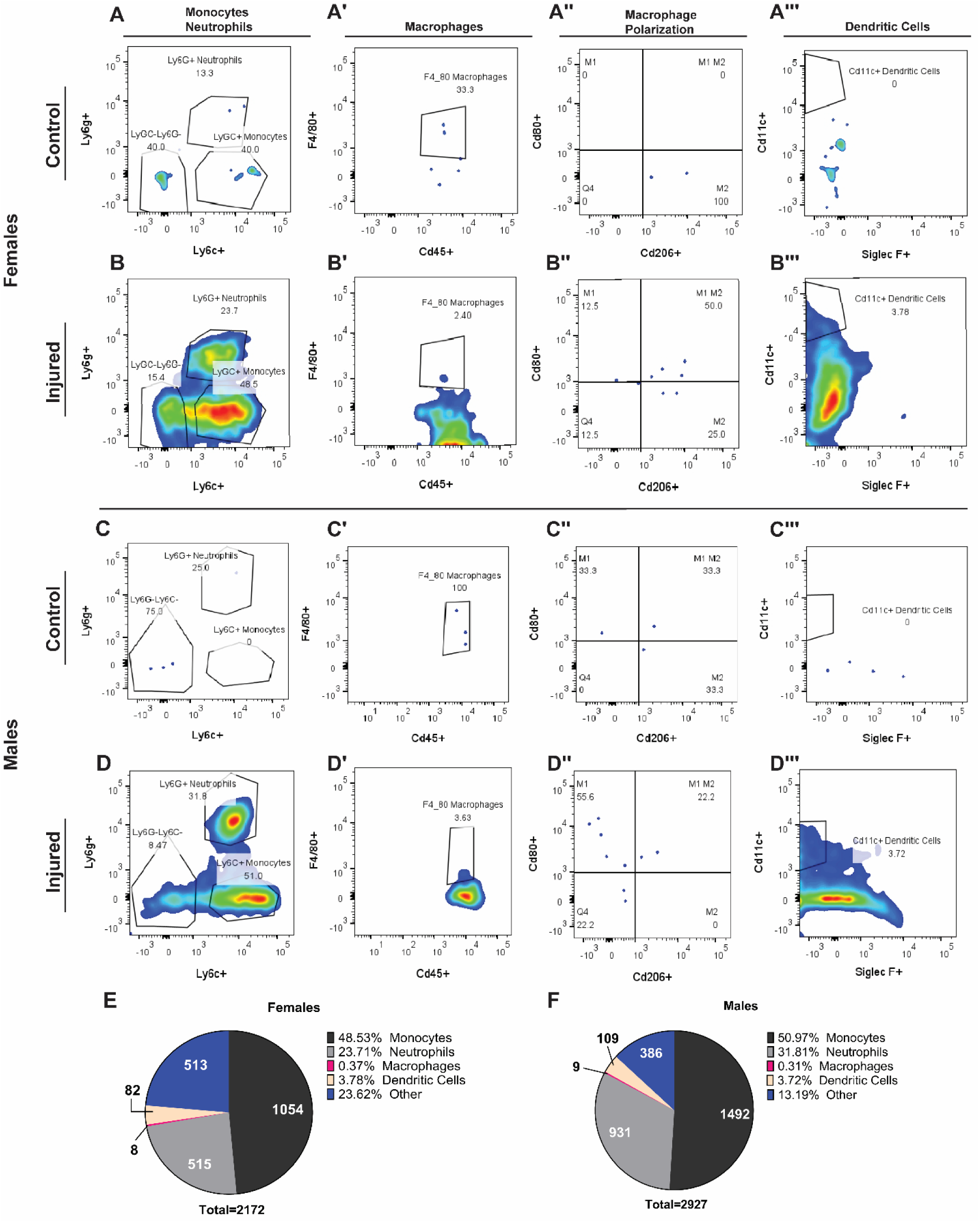
Males have higher percentages of responding Cd11b+ myeloid cells with injury at 3 dpi. Flow cytometry was used to identify infiltrating monocytes, neutrophils, macrophages, and dendritic cells in female (A, B) and male (C, D) mice from Cd45+Cd11b+ immune cells at 3dpi. (B) Injured female and (D) Injured male IVDs have a similar increase in myeloid cell populations when compared to controls (A, C), though males have a slightly higher increase in the most abundant myeloid populations, monocytes, and neutrophils. The percentage of myeloid subtypes represented in (E) female injured IVDs is very similar to those in (F) males. The totals in panels E, F are the total number of Cd11b+ myeloid cells injured samples. Chi-squared analysis of the difference in the proportions of female to male myeloid subtypes: χ2(4, N = 10) = 107.4, p = 0.0001.

F4/80+ macrophages were identified by sub-gating from the Ly6g-Ly6c-cell population and represented a very small population of Cd11b+ cells in control (**Figure 3A’, C’)** and injured tissues in both sexes (**Figure 3B’, D’**). We assessed macrophage polarization by using Cd206 to identify M2 polarized macrophages and Cd80 to identify M1 polarized macrophages. M2 macrophages are classically considered anti-inflammatory, and M1 are considered proinflammatory^27^. In females, 100% of the resident F4/80+ macrophages are M2 polarized (**Figure 3A’’**), but the majority are double positive for both M1 and M2 cell markers or M2 only (**Figure 3B’’**). Males have an even split between M1 and M2 polarized resident macrophages (**Figure 3C’’**), but most macrophages become M1 polarized with injury (**Figure 3D’’**).

We also determined changes in dendritic cells, another important APC in repair^26^. There are no resident Cd11c+ dendritic cells in either sex (**Figure 3A’’’, C’’’**), but both sexes have over a 3% increase with injury (**Figure 3B’’’, D’’’**). Though not as commonly associated with repair than macrophages, neutrophils or dendritic cells, the rapid recruitment of eosinophils into injured muscle has been shown to stimulate tissue repair^28^. Therefore, we also measured the response of Siglec F+ eosinophils in the IVD injury response. Eosinophils comprise a very small percentage of Cd11b+ cells in both sexes have respond similarly to injury in both sexes (**Figure S2**). Comparative analysis of the total cell numbers and percentages of the myeloid subtypes in injured IVDs of both sexes show subtle differences in the proportions of each subtype in females (**Figure 3E**) and males (**Figure 3F**), where males have higher percentages and a higher total number of infiltrating cells.

### Cd4-Cd8-T cells (DN T) dominate the lymphocyte response during acute injury in females but not males

Next, we identified the different subtypes of lymphocyte subtypes due to injury at 3 dpi in Cd45+ immune cells. The largest percentage of resident immune cells are Cd3+ T cells in control females, but not in males (**Figure 4A, C**). With injury, the highest responding cell types are myeloid cells in both sexes, but males have over 10% less infiltrating Cd3+ T cells in response to injury when compared to females (**Figure 4B, D**). We the sub-gated the Cd3+ cell population to identify the subtypes of responding T cells the and discovered that the overwhelming majority of resident T cells in control female mice are Cd4-Cd8-double negative T cells (DN T cells) with a small percentage being Cd8+ T cells, while in males the resident lymphocytes consist of a low percentage of DN T cells and a larger percentage of Cd8+ cells than females (**Figure 4A’, B’**). This trend persists with injury where DN T cells present over 96% of the population of T cells with less than 3% of Cd8+ T cells, but injured male mice have 80% DN T cells and 20% Cd8+ T cells (**Figure 4B’, D’**).

**Figure 4:**
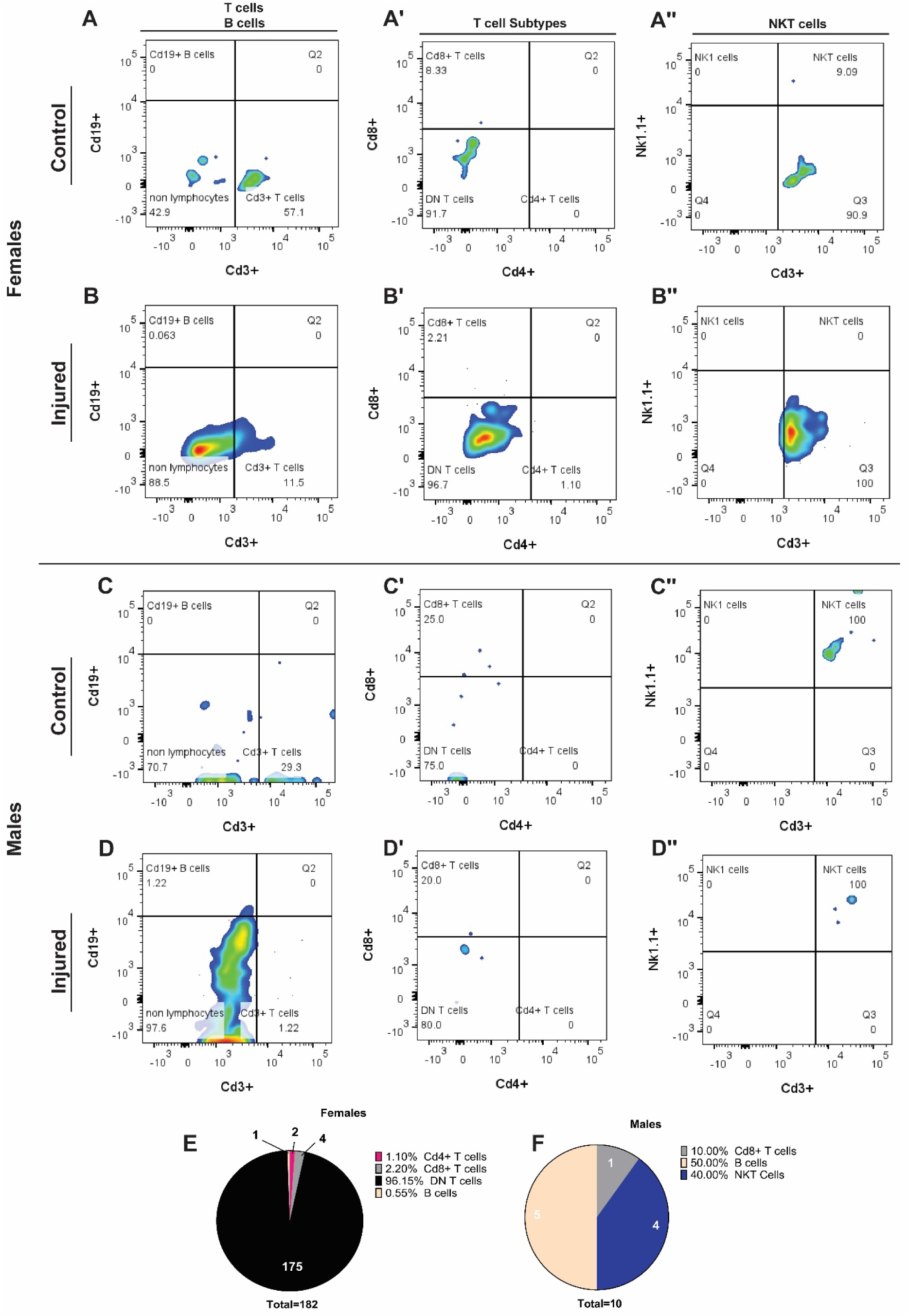
Cd4-Cd8-T cells (DN T) dominate the lymphocyte response during acute injury in females but not males. Flow cytometry used to identify infiltrating T cell subtypes, B cells and NKT cells in female (A, B) and male (C, D) mice from Cd45+ immune cells at 3dpi. (B) Injured female and (D) Injured male IVDs have a shift in the proportions of the lymphocytes subtypes when compared to controls, especially in the percentage of female T cell subtypes with injury (B’, D’). The lymphocyte response in (E) female injured IVDs is drastically different from those in (F) males where the female response is dominated by DN (Cd4-Cd8-Double Negative) T cells and the male response is dominated by Cd8+ cytotoxic T cells and B cells. The totals in panels E, F are the total number of lymphocytes injured samples. Chi-squared analysis of the difference in the proportions of female to male lymphocyte subtypes: χ2(4, N = 10) = 158.9, p = 0.0001.

NK T cells a very small portion of T lymphocytes, 0.1-1%, that express both Nk1.1 and Cd3 but do not have to express Cd4 or Cd8 markers^29^. To determine if the DN T cell population contains any NK1.1 expressing NK T cells, we sub-gated from the DN T cell population in females (**Figure 4A’’, B’’**) and males (**Figure 4C’’, D’’**). There is a very small population of Cd4-Cd8-Nk1.1+ NK T cells in control female IVDs (**Figure 4A’’**) but none with injury (**Figure 4B’’**). In males, 100% percent of the DN T cell population are NK T cells in both control (**Figure 4C’’**) and injured IVDs (**Figure 4D’’**). DN T cells usually represent a small portion of circulating T cells, 1-3%, and have been shown to have similar functions as regulatory T cells (Tregs)^30^. Of the 182 infiltrating lymphocytes in injured female mice, over 175 are DN T cells and the rest are smaller proportions of Cd4+ T cells, Cd8+ T cells and B cells (**Figure 4E**). In contrast, there is much smaller number of infiltrating lymphocytes (10) in males, and most of those cells are B cells and NKT cells with a smaller percentage of Cd8+ T cells (**Figure 4F**).

### Males exhibit a reduction in the proportions of Cd11b+ myeloid subtypes at 7 dpi despite having peak *Cd11b* gene regulation

Longitudinal analyses of infiltrating immune cells in response to injury is an important assessment to characterize how IVD-immune cell crosstalk changes over the course of the acute injury response. *Cd11b* has the greatest gene expression in the male injury response at 7 dpi (**Figure 1F**). To determine longitudinal changes in infiltrating Cd11b+ myeloid cells in males, we identified myeloid cell subtypes at 7 dpi and compared these data to observations from a more acute injury response time point, 3 dpi. At 7 dpi, the number of resident Cd11b+ myeloid cells is very small with the majority comprised of neutrophils and Ly6g-Ly6c-cells, similar to the response at 3 dpi (**Figure 5A, A’, Figure 3A**). Ly6g-Ly6c-cells are the largest responding population in both control and sub-gating this population to identify macrophages showed that only a small percentage are F4/80+ macrophages (**Figure 5B, B’**). Macrophage polarization shifts from being a split between M2 and double positive for M1/ M2 markers in controls to and increased population of M1 macrophages with injury (**Figure 5A’’, B’’**). For Cd11c+ dendritic cells, there are no resident cells in controls, but they represent 4.22% in inured IVDs, a percentage slightly higher that that seen at 3 dpi (**Figure 5A’’’, B’’’, Figure 3A’’’, B’’’**). Overall, the highest infiltrating population of Cd11b+ myeloid cell at 7 dpi is monocytes (**Figure 5C**). Though this observation is true for 3 dpi as well, there are differences in the proportions of the myeloid cell types between the two time points. Monocytes and neutrophils represent a smaller percentage while macrophages, dendritic cells have increased percentages at 7 dpi when compared to 3 dpi (**Figure 5C**, **Figure 3F**). The total number of infiltrating myeloid cells in reduced (829) compared to 3 dpi (**Figure 5C**, **Figure 3F**), but there is an increase in the number of responding macrophages and neutrophils with injury (**Figure 5D**).

**Figure 5:**
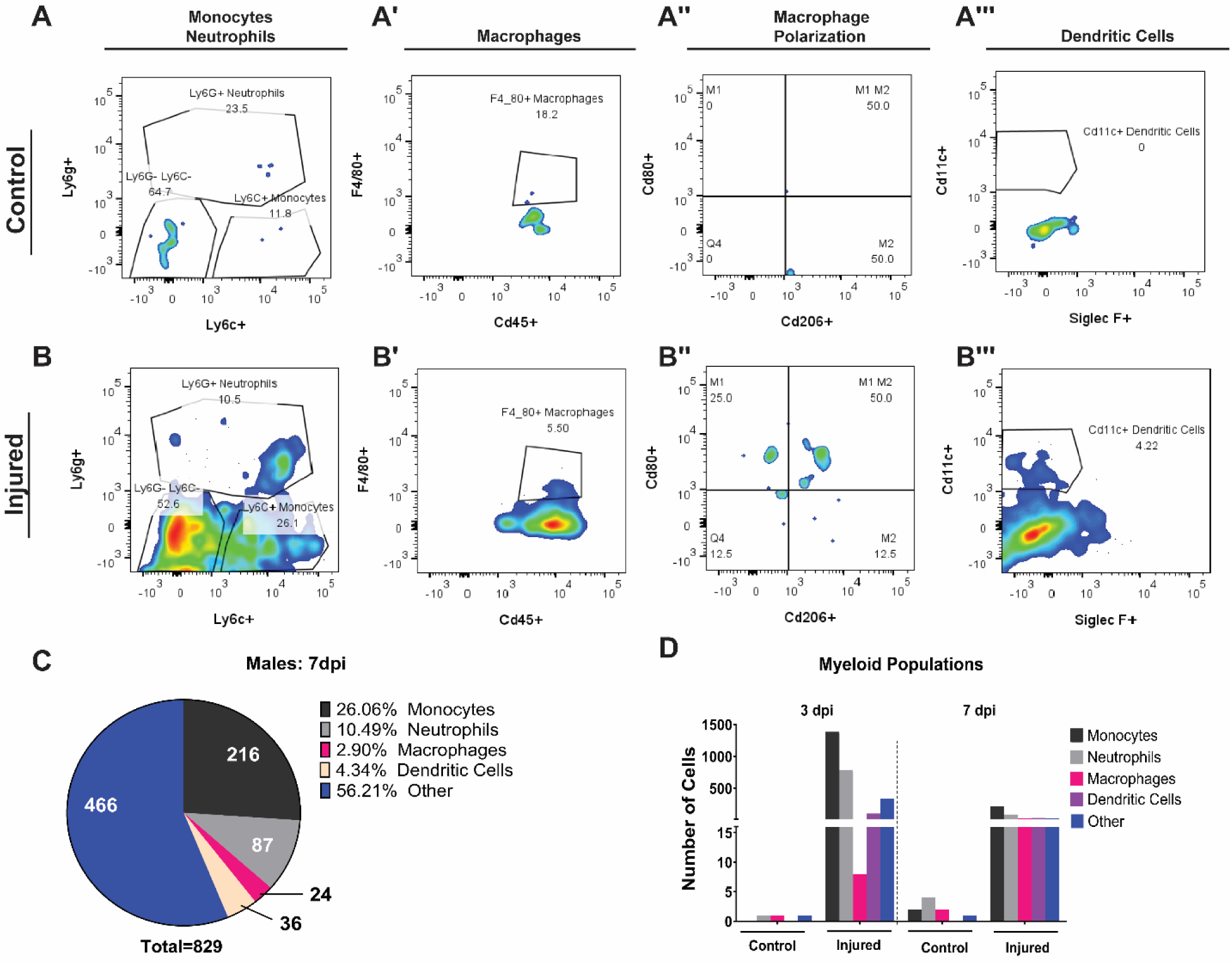
Males exhibit a reduction in the proportions of Cd11b+ myeloid subtypes at 7 dpi despite having peak *Cd11b* gene regulation. Changes in the subtypes of responding Cd11b+ myeloid cells at 7 dpi in response to injury compared to 3 dpi was measured using flow cytometry. There is a shift in the proportions of the responding myeloid cell subtypes in (A) Control compared to (B) Injured IVDs in males at 7 dpi that varies dependent upon the cell type. (C) The total number of cells has decreased and the proportions of the subtypes that respond at 7 dpi when compared to 3dpi has altered. (D) The comparison of myeloid subtype cell numbers between 3 dpi and 7 dpi show that the myeloid cell response is overall blunted at 7dpi, but more macrophages respond. Chi-squared analysis of the difference in the proportions of female to male lymphocyte subtypes: χ2(4, N = 10) = 773.6, p = 0.0001.

### 19 dpi marks peak *Cd3 gene* expression for T cells and concomitant reduction of DN T cells in females

Cd3+ T cells are the most abundant responding lymphocytes in females and their peak time of gene regulation is 19 dpi (**Figure 1F**, **Figure 4A, B**). Therefore, we used flow cytometry to identify responding lymphocytes cell types at 19 dpi and compared the changes in lymphocyte populations to a more acute time point, 3 dpi. At 19 dpi, there is an increase in the percentage of infiltrating Cd3+ T cells with injury when compared to controls (**Figure 6A, B**), though this percentage is reduced from that at 3 dpi (**Figure 4B**). Unlike 3 dpi where the vast majority of responding T cells are DN T cells in control and injured IVDs (**Figure 4A’, B’**), at 19 dpi the T cell populations in controls are nearly evenly split between Cd8+ and DN T cells while injured IVDs have majorly Cd8 + T cells (**Figure 6A’, B’**). NK1.1 expressing NK T cells were sub-gated from the DN T cell population and none of the DN T cells are NK T cells (**Figure 6A’’, B’’**).

**Figure 6:**
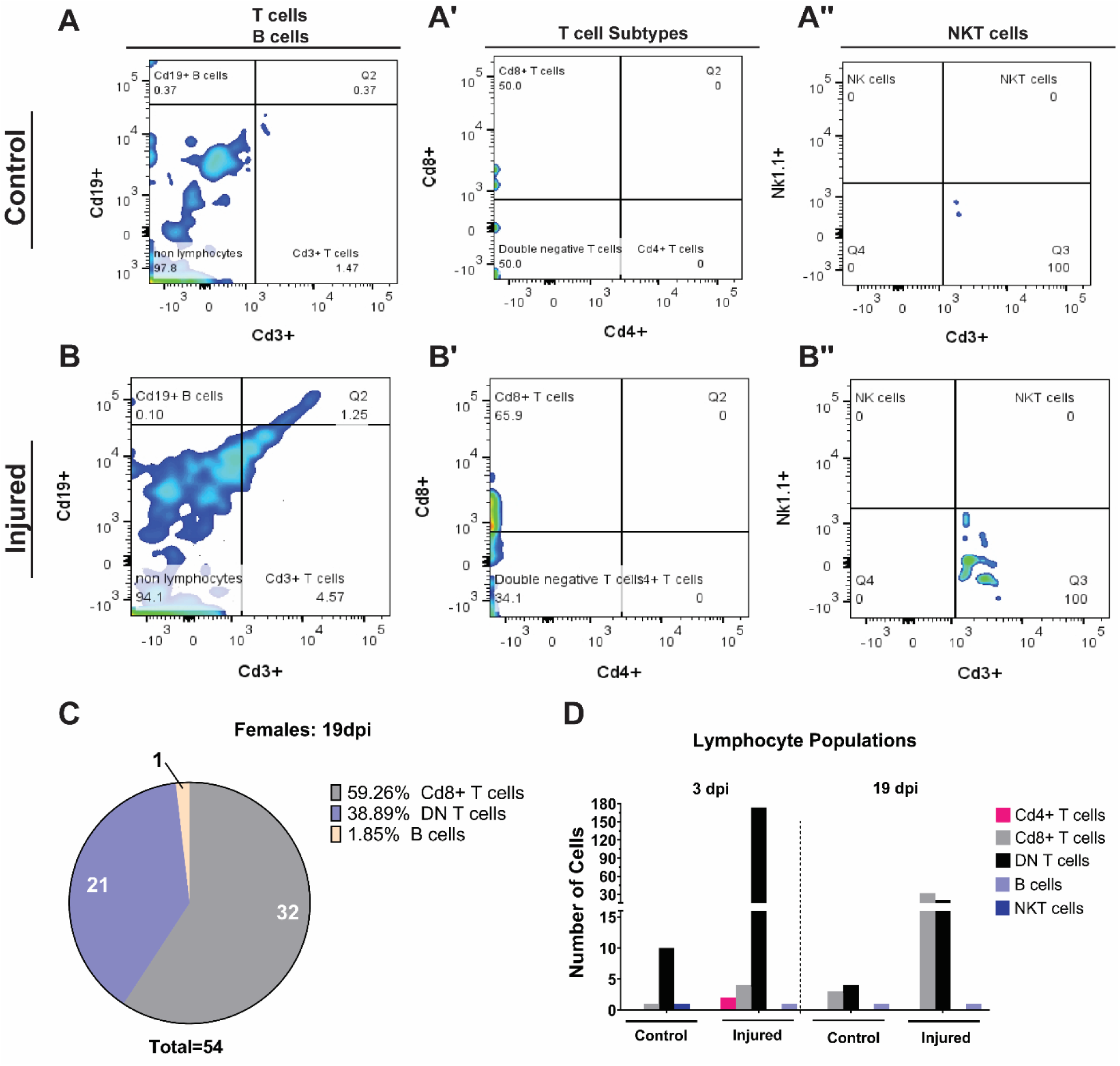
19 dpi marks peak *Cd3 gene* expression for T cells and concomitant reduction of DN T cells in females. Changes in the subtypes of responding lymphocytes at 19 dpi in response to in injury when compared to 3 dpi was measured using flow cytometry. There is an increase in infiltrating Cd3+ T cells when comparing (A) Control and (B) Injured IVDs. (C)There is a decrease and shift in the proportion of the lymphocyte subtypes at 19 dpi from 3dpi proportions where Cd8+ cells dominate at 19 dpi. (D) The comparison of lymphocyte subtype cell numbers show the lymphocyte response is dominated by DN T cells at 3 dpi and Cd8+ t cells at 19dpi with fewer total cell numbers. Chi-squared analysis of the difference in the proportions of female to male lymphocyte subtypes: χ2(3, N = 8) =90.105.4, p = 0.0001.

Since Cd3+ T cell gene expression oscillates and the lymphocytes that return at 19 dpi could be naïve or memory T cells, we sub-gated the Cd8+ and DN T cell populations with Cd44 and Cd62L markers to determine the presence of Cd44+ memory, Cd62+ naïve, or Cd44+Cd62L+ central memory T cells. Responding Cd8+ T cells are majorly naïve or central memory cells with injury in contrast to DN T cells that are majorly effectors (**Figure S3**). The analysis of the proportions and types of responding lymphocytes at 19 dpi with injury revealed the total number of lymphocytes is smaller, 54, and the proportions of responding lymphocytes at 19 dpi, has shifted to a Cd8+ T cell response (**Figure 6C**). Comparisons of the number of infiltrating cells for each lymphocyte subtype show a pivot in they the type and number of lymphocytes with injury and over time where this is an increased percentage of Cd8+ T cells and a reduction in DN T cells (**Figure 6D**).

### γδ T cells localize to the IVD in females after injury

To determine the localization of infiltrating immune cells, we used immunofluorescence to visualize the presence of Cd45, Cd11b, Cd3, or TCR gamma delta (TCRγδ) positive cells within the annulus fibrosus and nucleus pulposus. TCRγδ is the T cell receptor expressed on Cd3+Cd4-Cd8-T cells (DN T cells) and have been shown to have anti-inflammatory, pro-repair functions after injury^31,32^. There are a small number of resident Cd45+ and Cd11b+ immune cells within Control IVDs that are mostly restricted to the nucleus pulposus (**Figure 7A, B, Figure S4**). With injury, there is an increase in Cd45 and Cd11b expressing cells in both sexes in the IVD space, particularly the nucleus pulposus, and in the neighboring tissues (**Figure 7A’, B’, C, D, Figure S5**).

**Figure 7:**
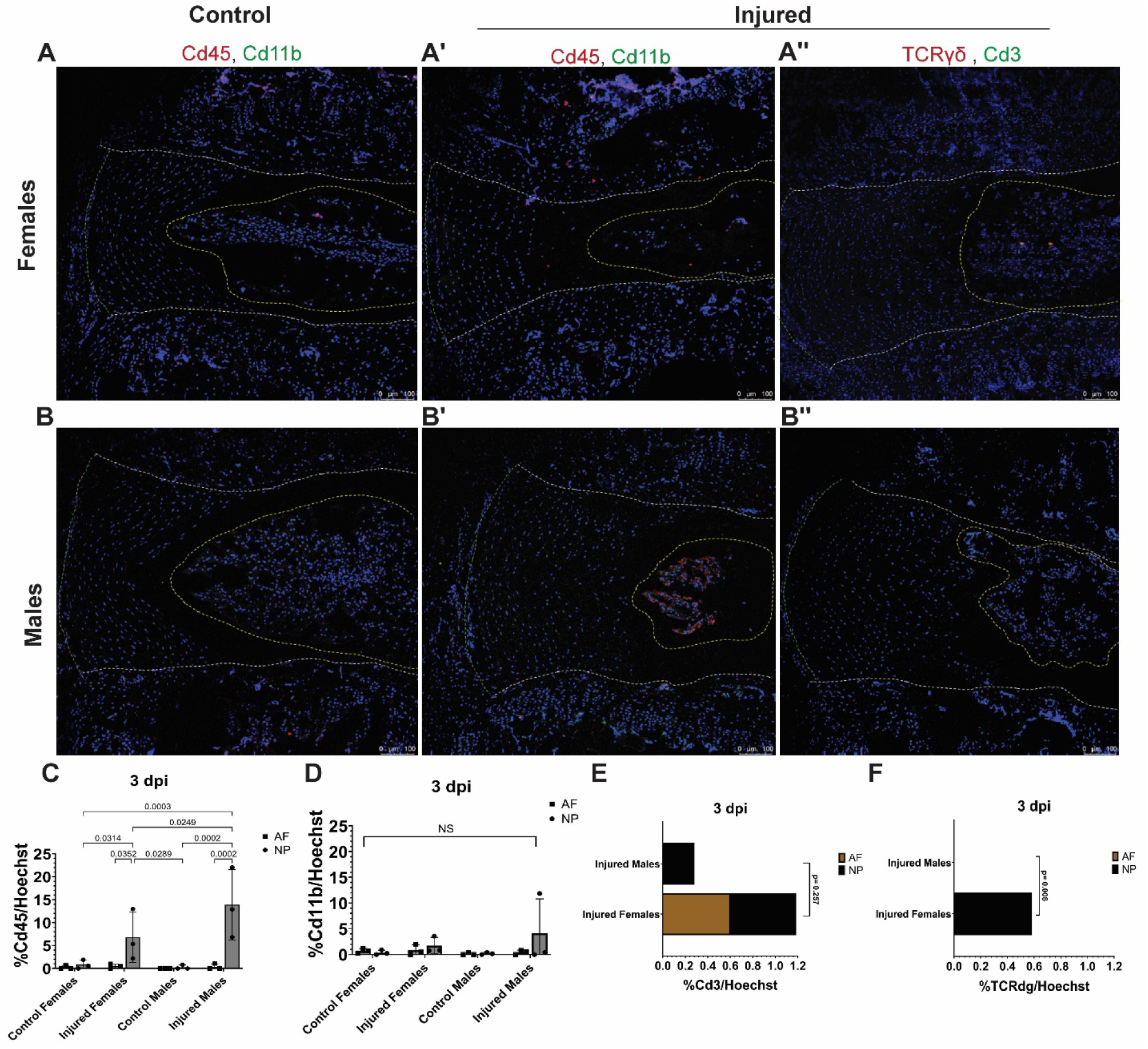
γδ T cells localize to the IVD in females after Injury. Localization of Cd45+/Cd11b+ or Cd3+/TCRγδ+ expressing cells was analyzed with immunofluorescence. Cd45+ and Cd11b+ cells in Control (A) female and (B) male samples and Injured (A’) female and (B’) male samples were identified. Cd3+ and TCRγδ+ cells in Injured (A’’) female and (B’’) male samples were identified as well. Quantification of the percentage of (C) Cd45+, (D) Cd11b+, (E) Cd3+ and (F) TCRγδ+ cells divided by total Hoechst positive nuclei in the annulus fibrosus and nucleus pulposus was analyzed.

Next, we wanted to determine the localization of Cd3+ T, the highest responding lymphocyte in response to injury (**Figure 4A-D**), and TCRγδ + T cells to determine if the DN T cells identified with flow cytometry were these specialized T cells. We identified the presence of infiltrating Cd3+ T cells that are TCRγδ+ only in female injured IVDs (**Figure 7A’’, B’’, E, F**). There were also more Cd3+ T cells that infiltrate females than males and no presence of TCRγδ cells in males, corroborating our flow cytometry data (**Figure 7E, F**).

## Discussion

This study is the first to conduct a comprehensive, longitudinal analysis of the IVD acute injury response that identifies the types, temporal regulation, and sex-related divergences of responding immune cells. By using a combinatorial approach which employed analysis of longitudinal changes in gene regulation, IVD tissue morphology, the dynamics of infiltrating immune cells and their tissue localization in response to injury, we were able to characterize sex-specific differences in the response of myeloid and lymphocyte cell populations and identify the involvement of previously unreported immune cell types like γδ and NK T cells. These findings can be used to exploit approaches to modulate immune cell subsets to increase the repair capacity of the IVD, a tissue with extremely limited regenerative capability^24^.

A major finding from this study is the sex-related divergences in immune cell gene regulation with injury. The longitudinal expression patterns of the immune cell and angiogenesis genes permitted the temporal mapping of the IVD acute versus chronic injury response and the approximation of the time windows of the three tissue repair stages based on the alternating patterns of gene regulation between the immune cells and angiogenic factors. These data aided in the identification of key regulatory time points throughout the injury response, 3,7,12,19 and 42 dpi, and pinpoint differences in gene regulation between sexes (**Figure 1**). Analysis of the longitudinal changes in injured tissue morphology determined progressive degeneration and the lack of repair independent of sex. Though the amount of degeneration is similar between sexes early in the acute injury response at 3 dpi, males appear to have accelerated degeneration throughout the acute response until 42 dpi (**Figure 2)**. This indicates that 12 week old females are slightly protected from degenerative changes. This finding is supported in humans since young female patients have less IVD degeneration than males though this trend reverses with aging^33,34^. Rodent studies have also determined sex-specific differences where male rats have IVD injury-associated pain 6 weeks after injury, but females do not^35^. Though others have reported no detectable changes in histology at 1 week or 6 weeks post injury between sexes, differences in the method of injury must be considered. Our method is a severe, traumatic injury where a bilateral needle puncture was strategically used to invoke a robust immune response while other studies used a more conservative unilateral injury^35–37^. Our previous work has shown that injuries that transverse the full width of the IVD (bilateral) have more severe degeneration than those that transverse only the partial width of the IVD (unilateral), highlighting the differential outcomes that come with injury severity^38^.

There are a number of studies that documented immune cell presence in herniated or degenerate IVDs, while few examine the cascade acutely following injury^39–41^. Macrophages have been shown to be present in injured or herniated IVD tissue in rodent and human studies at acute time points post injury, but none have identified diverse myeloid cell types, their longitudinal response, and the differences between sexes as shown in this study^40–44^. Analysis of the responding myeloid cells determined monocytes and neutrophils as the most abundant responding cell type in at 3 dpi with injury, and there is a difference in the proportions of monocytes, neutrophils, macrophages, dendritic cells, and eosinophils between sexes (**Figure 3**). The reduction of responding myeloid cells in males at 7 dpi is accompanied by an increase in the proportion and number of macrophages only when compared to 3 dpi. Macrophage polarization also shifted from M1 at 3 dpi to a M1 and M1/M2 polarized macrophages at 7 dpi (**Figure 5**). as well, but further studies into the implication of this polarization shift needs to be conducted.

The identities of lymphocytes during IVD acute injury is less examined and usually measured through surrogates such as cytokine production or transcriptomic changes. Human patient discectomy samples have been shown to contain Cd4+ T cells, but these tissues are in a chronic state of IVD degeneration and do not give insight into the role the T cells play in the acute healing stages^43,45^. Rodent models have shown the presence of Cd4+ or Cd8+ T cells at isolated time points, but rarely address sex-related differences or subtype differences^45–47^. We discovered drastic differences in the responding lymphocytes between the sexes at 3 dpi where DN T cells dominated the female response but Cd8+ and NK T cells were responders in males with injury (**Figure 4**). The prevalence of these T reg like DN T cells in females could be associated with the reduced degenerative changes that injured females have when compared to males. In contrast, the response of Cd8+ and NK T cells in males, which have been shown to cause deleterious effects to tissue repair or contribute to cytotoxicity, could contribute to the greater degenerative changes seen via histology^16,48^. The longitudinal response in females at 19 dpi showed a shift in lymphocytes where Cd8+ T cells were the highest responders could contribute to why repair is still impaired in females (**Figure 6**). The large and specific presence of DN T cells in only females led us to discover the presence of Cd3+ TCRγδ+ T cells with immunofluorescence (**Figure 7**). To our knowledge, we are the first to discover the presence and sex-specific regulation of γδ T cells in response to IVD injury. γδ T cells could also contribute to the reduced degenerative changes seen in females since these cells are shown to promote healing in bone and muscle^49,50^.

The mechanisms that promote immune cell infiltration into the IVD due to injury were not investigated. Previous work has demonstrated that applying damaging mechanical loads promotes production of neutrophil-recruiting chemokines and mechanosensitive ion channel activation temporally regulates immune cell-recruiting chemokines^51^. Endogenous production of immune-related chemokines from IVD cells could partially explain how immune cells home to injured IVDs. A limitation of this study is the small number of cells available per mouse caudal IVD for flow cytometry analyses. Though we pooled 15 discs from 3 animals per flow cytometry run, the total number of cells collected is small, ca. 23,000. Despite the small cell numbers, we detected distinct immune cell populations, but additional pooling of tissues may be necessary for more in depth experiments. Another limitation is this study does not consider how the estrus cycle affects the immune system response. Though studies have shown that estrus cycle has no effect on IVD degeneration outcomes or resultant pain experienced in the rat model, those studies did not examine any effects on the responding immune cells^35^. This is important since studies have shown that immune cells can fluctuate over the estrus cycle^52^. Lastly, these findings are in young animals and the majority of degeneration occurs in aged patients. These findings need to be expanded to aged animals and human samples to assess changes in the immune response with aging and across species, which we will explore in future studies. These studies begin to unravel complex the sex-related differences in IVD-immune crosstalk and the effect on degeneration. The identification of novel immune cell populations, like NK T and γδ T cells, and their role in acute injury provides new targetable cell types to modulate the repair capacity of the IVD.

## Acknowledgements

This work was conducted with funding support from National Institute of Arthritis and Musculoskeletal and Skin Diseases: R01AR074441, R01AR077678, R21AR081517, P30 AR074992, T32 Postdoctoral Training in Regenerative Medicine (T32 EB028092) from the National Institute of Biomedical Imaging and Bioengineering, and the Rita Levi-Montalcini Postdoctoral Fellowship in Regenerative Medicine from the Center of Regenerative Medicine at Washington University. Thank you to Roberta Faccio’s lab members and the Flow Cytometry & Fluorescence Activated Cell Sorting Core for flow cytometry training and usage of the core’s analyzers. Thank you to Tang lab members, Hong Joo Moon and Christian Gonzalez, for assistance with degeneration scoring.

## Author Contributions

SWC and SYT designed the research study, contributed to data interpretation, and wrote the manuscript. SWC, REW, LM, and GWDE performed research and analyzed data. All co-authors reviewed and revised the manuscript.

## Competing Interests

The authors declare no competing interests.

## Data Availability

All data collected for this study is available upon request.

## Notes

### Competing Interest Statement

The authors have declared no competing interest.

## References

1. Tavakoli J, Diwan AD, Tipper JL. Advanced Strategies for the Regeneration of Lumbar Disc Annulus Fibrosus. Int J Mol Sci. Jul 10 2020;21(14)doi:10.3390/ijms21144889

2. Romereim SM, Johnston CA, Redwine AL, Wachs RA. Development of an in vitro intervertebral disc innervation model to screen neuroinhibitory biomaterials. J Orthop Res. May 2020;38(5):1016–1026. doi:10.1002/jor.24557

3. Wang SZ, Rui YF, Tan Q, Wang C. Enhancing intervertebral disc repair and regeneration through biology: platelet-rich plasma as an alternative strategy. Arthritis Res Ther. 2013;15(5):220. doi:10.1186/ar4353

4. Tang G, Zhou B, Li F, et al. Advances of Naturally Derived and Synthetic Hydrogels for Intervertebral Disk Regeneration. Front Bioeng Biotechnol. 2020;8:745. doi:10.3389/fbioe.2020.00745

5. Eming SA, Martin P, Tomic-Canic M. Wound repair and regeneration: mechanisms, signaling, and translation. Sci Transl Med. Dec 3 2014;6(265):265sr6. doi:10.1126/scitranslmed.3009337

6. Dou Y, Sun X, Ma X, Zhao X, Yang Q. Intervertebral Disk Degeneration: The Microenvironment and Tissue Engineering Strategies. Front Bioeng Biotechnol. 2021;9:592118. doi:10.3389/fbioe.2021.592118

7. Lyu FJ, Cui H, Pan H, et al. Painful intervertebral disc degeneration and inflammation: from laboratory evidence to clinical interventions. Bone Res. Jan 29 2021;9(1):7. doi:10.1038/s41413-020-00125-x

8. Abraham AC, Liu JW, Tang SY. Longitudinal changes in the structure and inflammatory response of the intervertebral disc due to stab injury in a murine organ culture model. J Orthop Res. Aug 2016;34(8):1431–8. doi:10.1002/jor.23325

9. Gonzalez AC, Costa TF, Andrade ZA, Medrado AR. Wound healing - A literature review. An Bras Dermatol. Sep-Oct 2016;91(5):614–620. doi:10.1590/abd1806-4841.20164741

10. Schultz GS, Chin GA, Moldawer L, Diegelmann RF. Principles of Wound Healing. In: Fitridge R, Thompson M, eds. Mechanisms of Vascular Disease: A Reference Book for Vascular Specialists. University of Adelaide Press © The Contributors 2011.; 2011.

11. Flint JH, Wade AM, Giuliani J, Rue J-P. Defining the Terms Acute and Chronic in Orthopaedic Sports Injuries:A Systematic Review. The American Journal of Sports Medicine. 2014;42(1):235–241. doi:10.1177/0363546513490656

12. Ono T, Takayanagi H. Osteoimmunology in Bone Fracture Healing. Curr Osteoporos Rep. Aug 2017;15(4):367–375. doi:10.1007/s11914-017-0381-0

13. Laumonier T, Menetrey J. Muscle injuries and strategies for improving their repair. J Exp Orthop. Dec 2016;3(1):15. doi:10.1186/s40634-016-0051-7

14. Walker J KS, Michelsons S, Creber K, Elliott C, Leask A, Hamilton D. Cell–matrix interactions governing skin repair: matricellular proteins as diverse modulators of cell function. Research and Reports in Biochemistry. 2015;5:73–88. doi:doi.org/10.2147/RRBC.S57407

15. Muire PJ, Mangum LH, Wenke JC. Time Course of Immune Response and Immunomodulation During Normal and Delayed Healing of Musculoskeletal Wounds. Front Immunol. 2020;11:1056. doi:10.3389/fimmu.2020.01056

16. Baht GS, Vi L, Alman BA. The Role of the Immune Cells in Fracture Healing. Curr Osteoporos Rep. Apr 2018;16(2):138–145. doi:10.1007/s11914-018-0423-2

17. Ziemkiewicz N, Hilliard G, Pullen NA, Garg K. The Role of Innate and Adaptive Immune Cells in Skeletal Muscle Regeneration. Int J Mol Sci. Mar 23 2021;22(6)doi:10.3390/ijms22063265

18. Larouche J, Sheoran S, Maruyama K, Martino MM. Immune Regulation of Skin Wound Healing: Mechanisms and Novel Therapeutic Targets. Adv Wound Care (New Rochelle). Jul 1 2018;7(7):209–231. doi:10.1089/wound.2017.0761

19. Crosio G, Huang AH. Innate and adaptive immune system cells implicated in tendon healing and disease. European cells & materials. Feb 18 2022;43:39–52. doi:10.22203/eCM.v043a05

20. Li M, Yin H, Yan Z, et al. The immune microenvironment in cartilage injury and repair. Acta Biomater. Mar 1 2022;140:23–42. doi:10.1016/j.actbio.2021.12.006

21. Shi Z, Yao C, Shui Y, Li S, Yan H. Research progress on the mechanism of angiogenesis in wound repair and regeneration. Front Physiol. 2023;14:1284981. doi:10.3389/fphys.2023.1284981

22. Zhang M, Fukushima Y, Nozaki K, et al. Enhancement of bone regeneration by coadministration of angiogenic and osteogenic factors using messenger RNA. Inflamm Regen. Jun 20 2023;43(1):32. doi:10.1186/s41232-023-00285-3

23. Liu X, Zhu B, Li Y, et al. The Role of Vascular Endothelial Growth Factor in Tendon Healing. Front Physiol. 2021;12:766080. doi:10.3389/fphys.2021.766080

24. Ju DG, Kanim LE, Bae HW. Intervertebral Disc Repair: Current Concepts. Global Spine J. Apr 2020;10(2 Suppl):130s–136s. doi:10.1177/2192568219872460

25. Melgoza IP, Chenna SS, Tessier S, et al. Development of a standardized histopathology scoring system using machine learning algorithms for intervertebral disc degeneration in the mouse model-An ORS spine section initiative. JOR Spine. Jun 2021;4(2):e1164. doi:10.1002/jsp2.1164

26. Shanley LC, Mahon OR, Kelly DJ, Dunne A. Harnessing the innate and adaptive immune system for tissue repair and regeneration: Considering more than macrophages. Acta Biomater. Oct 1 2021;133:208–221. doi:10.1016/j.actbio.2021.02.023

27. Murray PJ, Allen JE, Biswas SK, et al. Macrophage activation and polarization: nomenclature and experimental guidelines. Immunity. Jul 17 2014;41(1):14–20. doi:10.1016/j.immuni.2014.06.008

28. Heredia JE, Mukundan L, Chen FM, et al. Type 2 innate signals stimulate fibro/adipogenic progenitors to facilitate muscle regeneration. Cell. Apr 11 2013;153(2):376–88. doi:10.1016/j.cell.2013.02.053

29. Wu L, Van Kaer L. Natural killer T cells in health and disease. Front Biosci (Schol Ed). Jan 1 2011;3(1):236–51. doi:10.2741/s148

30. Wu Z, Zheng Y, Sheng J, et al. CD3(+)CD4(-)CD8(-) (Double-Negative) T Cells in Inflammation, Immune Disorders and Cancer. Front Immunol. 2022;13:816005. doi:10.3389/fimmu.2022.816005

31. Hu Y, Hu Q, Li Y, et al. γδ T cells: origin and fate, subsets, diseases and immunotherapy. Signal Transduct Target Ther. Nov 22 2023;8(1):434. doi:10.1038/s41392-023-01653-8

32. Dar HY, Perrien DS, Pal S, et al. Callus γδ T cells and microbe-induced intestinal Th17 cells improve fracture healing in mice. J Clin Invest. Apr 17 2023;133(8)doi:10.1172/jci166577

33. Takatalo J, Karppinen J, Niinimäki J, et al. Prevalence of degenerative imaging findings in lumbar magnetic resonance imaging among young adults. Spine. Jul 15 2009;34(16):1716–21. doi:10.1097/BRS.0b013e3181ac5fec

34. Teraguchi M, Yoshimura N, Hashizume H, et al. Prevalence and distribution of intervertebral disc degeneration over the entire spine in a population-based cohort: the Wakayama Spine Study. Osteoarthritis and cartilage. Jan 2014;22(1):104–10. doi:10.1016/j.joca.2013.10.019

35. Mosley GE, Wang M, Nasser P, et al. Males and females exhibit distinct relationships between intervertebral disc degeneration and pain in a rat model. Scientific reports. Sep 15 2020;10(1):15120. doi:10.1038/s41598-020-72081-9

36. Mosley GE, Hoy RC, Nasser P, et al. Sex Differences in Rat Intervertebral Disc Structure and Function Following Annular Puncture Injury. Spine. Sep 2019;44(18):1257–1269. doi:10.1097/brs.0000000000003055

37. Brent JM, Tian Z, Shofer FS, et al. Influence of Genetic Background and Sex on Gene Expression in the Mouse (Mus musculus) Tail in a Model of Intervertebral Disc Injury. Comp Med. Apr 1 2020;70(2):131–139. doi:10.30802/aalas-cm-19-000034

38. Walk RE, Moon HJ, Tang SY, Gupta MC. Contrast-enhanced microCT evaluation of degeneration following partial and full width injuries to the mouse lumbar intervertebral disc. Scientific reports. Sep 16 2022;12(1):15555. doi:10.1038/s41598-022-19487-9

39. Wang L, He T, Liu J, et al. Revealing the Immune Infiltration Landscape and Identifying Diagnostic Biomarkers for Lumbar Disc Herniation. Front Immunol. 2021;12:666355. doi:10.3389/fimmu.2021.666355

40. Virri J, Grönblad M, Seitsalo S, Habtemariam A, Kääpä E, Karaharju E. Comparison of the prevalence of inflammatory cells in subtypes of disc herniations and associations with straight leg raising. Spine. Nov 1 2001;26(21):2311–5. doi:10.1097/00007632-200111010-00004

41. Lee S, Millecamps M, Foster DZ, Stone LS. Long-term histological analysis of innervation and macrophage infiltration in a mouse model of intervertebral disc injury-induced low back pain. J Orthop Res. Jun 2020;38(6):1238–1247. doi:10.1002/jor.24560

42. Xiao L, Matharoo J, Chi J, et al. Transient depletion of macrophages alters local inflammatory response at the site of disc herniation in a transgenic mouse model. Osteoarthritis and cartilage. Jul 2023;31(7):894–907. doi:10.1016/j.joca.2023.01.574

43. Shamji MF, Setton LA, Jarvis W, et al. Proinflammatory cytokine expression profile in degenerated and herniated human intervertebral disc tissues. Arthritis Rheum. Jul 2010;62(7):1974–82. doi:10.1002/art.27444

44. Nakawaki M, Uchida K, Miyagi M, et al. Changes in Nerve Growth Factor Expression and Macrophage Phenotype Following Intervertebral Disc Injury in Mice. J Orthop Res. Aug 2019;37(8):1798–1804. doi:10.1002/jor.24308

45. Yao Y, Xue H, Chen X, et al. Polarization of Helper T Lymphocytes Maybe Involved in the Pathogenesis of Lumbar Disc Herniation. Iran J Allergy Asthma Immunol. Aug 2017;16(4):347–357.

46. Gorth DJ, Shapiro IM, Risbud MV. Transgenic mice overexpressing human TNF-α experience early onset spontaneous intervertebral disc herniation in the absence of overt degeneration. Cell Death Dis. Dec 18 2018;10(1):7. doi:10.1038/s41419-018-1246-x

47. Rohanifar M, Clayton SW, Easson GWD, et al. Single Cell RNA-Sequence Analyses Reveal Uniquely Expressed Genes and Heterogeneous Immune Cell Involvement in the Rat Model of Intervertebral Disc Degeneration. Applied Sciences. 2022;12(16):8244.

48. Schlundt C, Reinke S, Geissler S, et al. Individual Effector/Regulator T Cell Ratios Impact Bone Regeneration. Front Immunol. 2019;10:1954. doi:10.3389/fimmu.2019.01954

49. Mann AO, Hanna BS, Muñoz-Rojas AR, et al. IL-17A-producing γδT cells promote muscle regeneration in a microbiota-dependent manner. J Exp Med. May 2 2022;219(5)doi:10.1084/jem.20211504

50. Ono T, Okamoto K, Nakashima T, et al. IL-17-producing γδ T cells enhance bone regeneration. Nature communications. Mar 11 2016;7:10928. doi:10.1038/ncomms10928

51. Easson GWD, Savadipour A, Gonzalez C, Guilak F, Tang SY. TRPV4 differentially controls inflammatory cytokine networks during static and dynamic compression of the intervertebral disc. JOR Spine. Dec 2023;6(4):e1282. doi:10.1002/jsp2.1282

52. Oertelt-Prigione S. Immunology and the menstrual cycle. Autoimmun Rev. May 2012;11(6-7):A486-92. doi:10.1016/j.autrev.2011.11.023

