## Supplementary Figures and Tables for "Analysis of Infiltrating Immune Cells Following Intervertebral Disc Injury Reveals Recruitment of Gamma-Delta (γδ) T cells in Female Mice"

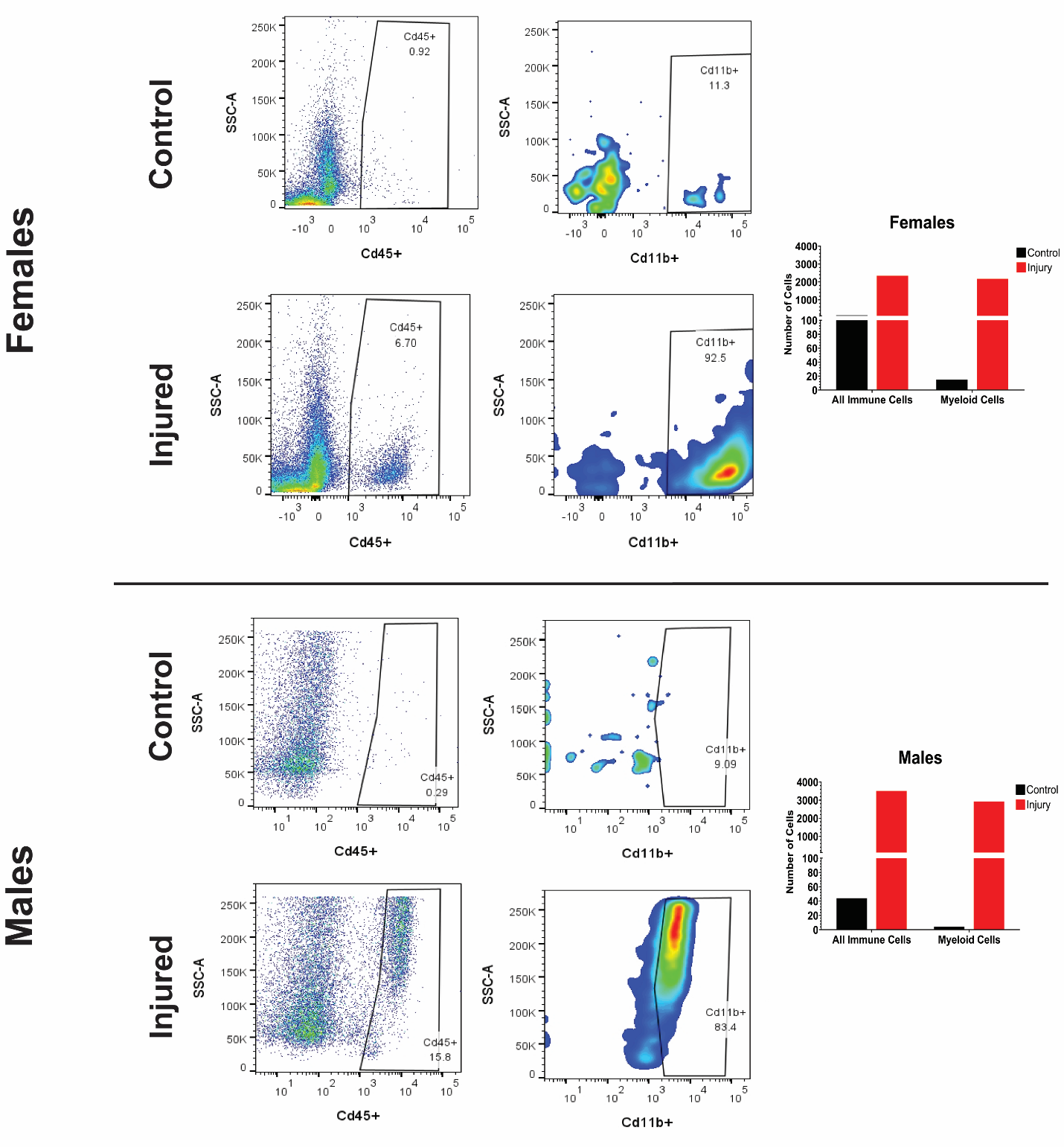
**Figure S1: Analysis of responding Cd45+ and Cd11b+ immune cells with injury.** Flow cytometry was used to quantify Cd45+ cells and identify how many of the Cd45+ cells are Cd11b+ myeloid cells in control and injured IVDs in both sexes.

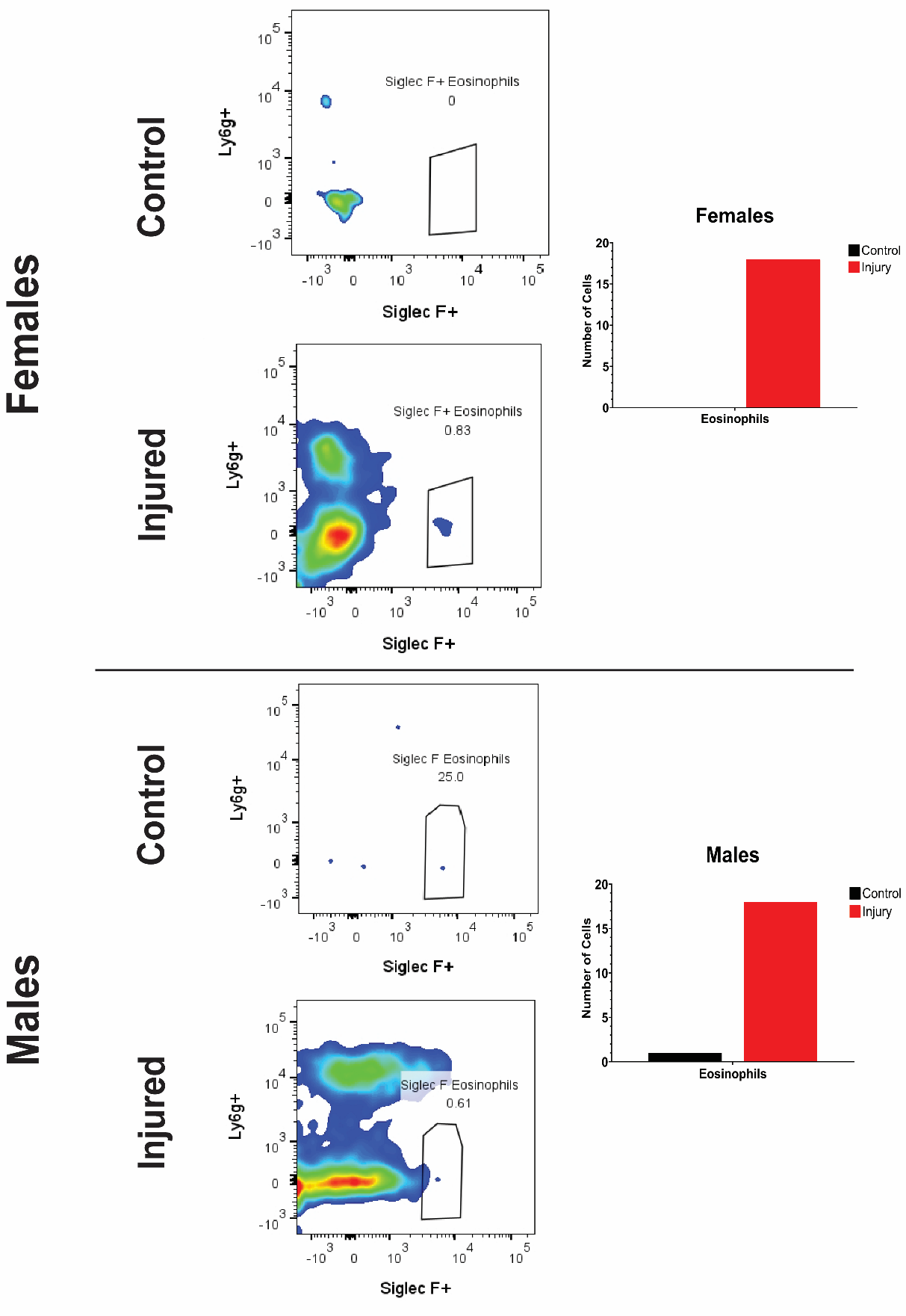

**Figure S2: Siglec F+ eosinophils are a small percentage of responding Cd11b+ myeloid cells.** Females had no eosinophils present in controls and a small number of infiltrating eosinophils with injury. Males had one detectable eosinophil in controls and an increase in eosinophil number with injury.

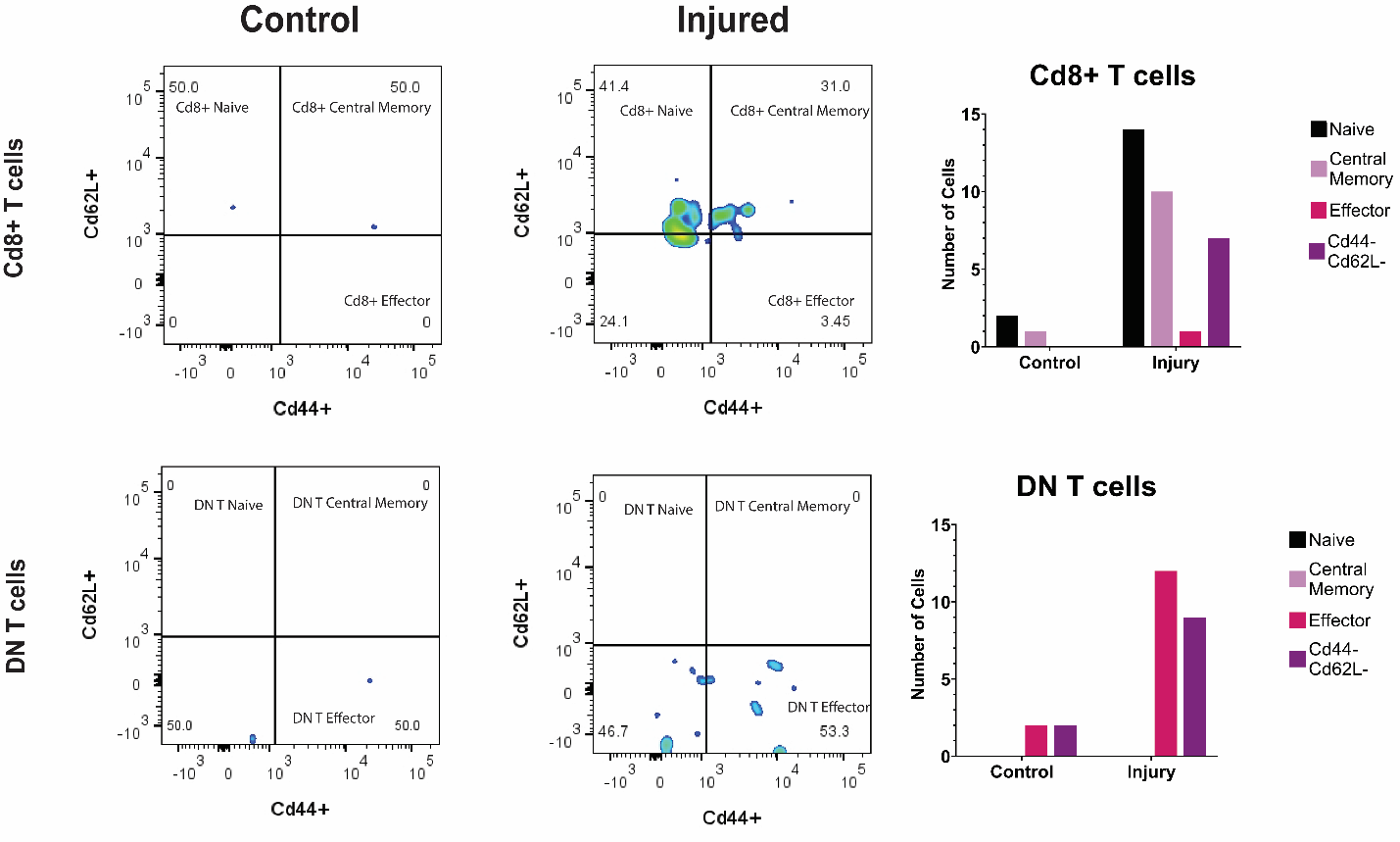

**Figure S3: Identification of the naïve, central memory, and effector Cd8+ and DN T cells at 19 dpi in females.** Cd62L+ naïve, Cd44 effector, and Cd62L+Cd44+ central memory Cd8 and DN T cells were identified and quantified in female control and injured IVDs at 19 dpi.

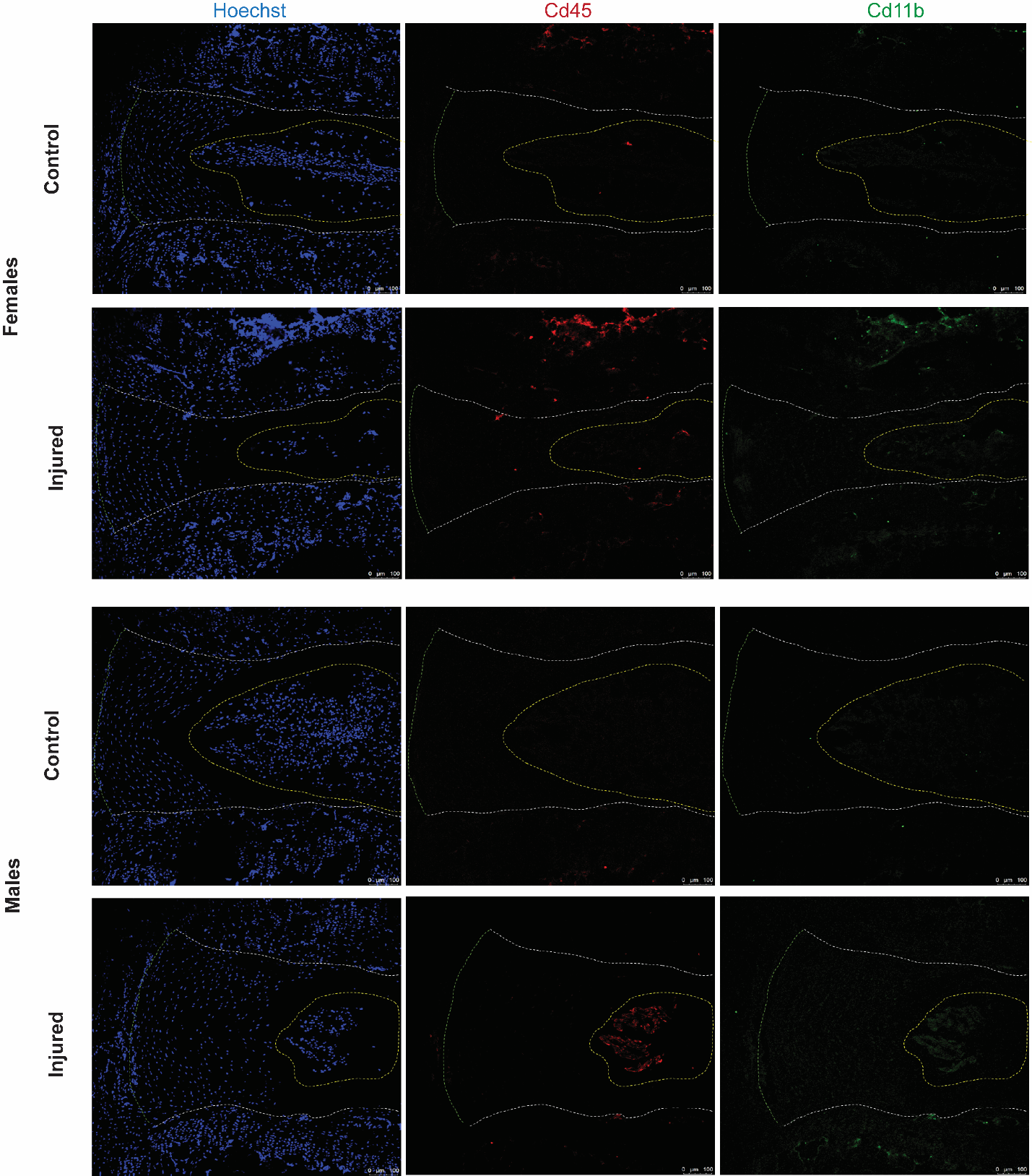

**Figure S4:** **Single channel analysis of Cd45 and Cd11b immunofluorescence.** Separated Hoechst (nuclei), Cd45, and Cd11b immunofluorescence channels from control and injured IVDs in both sexes permit the better visualization of the localization of immune cells within and at the periphery of IVD tissue.

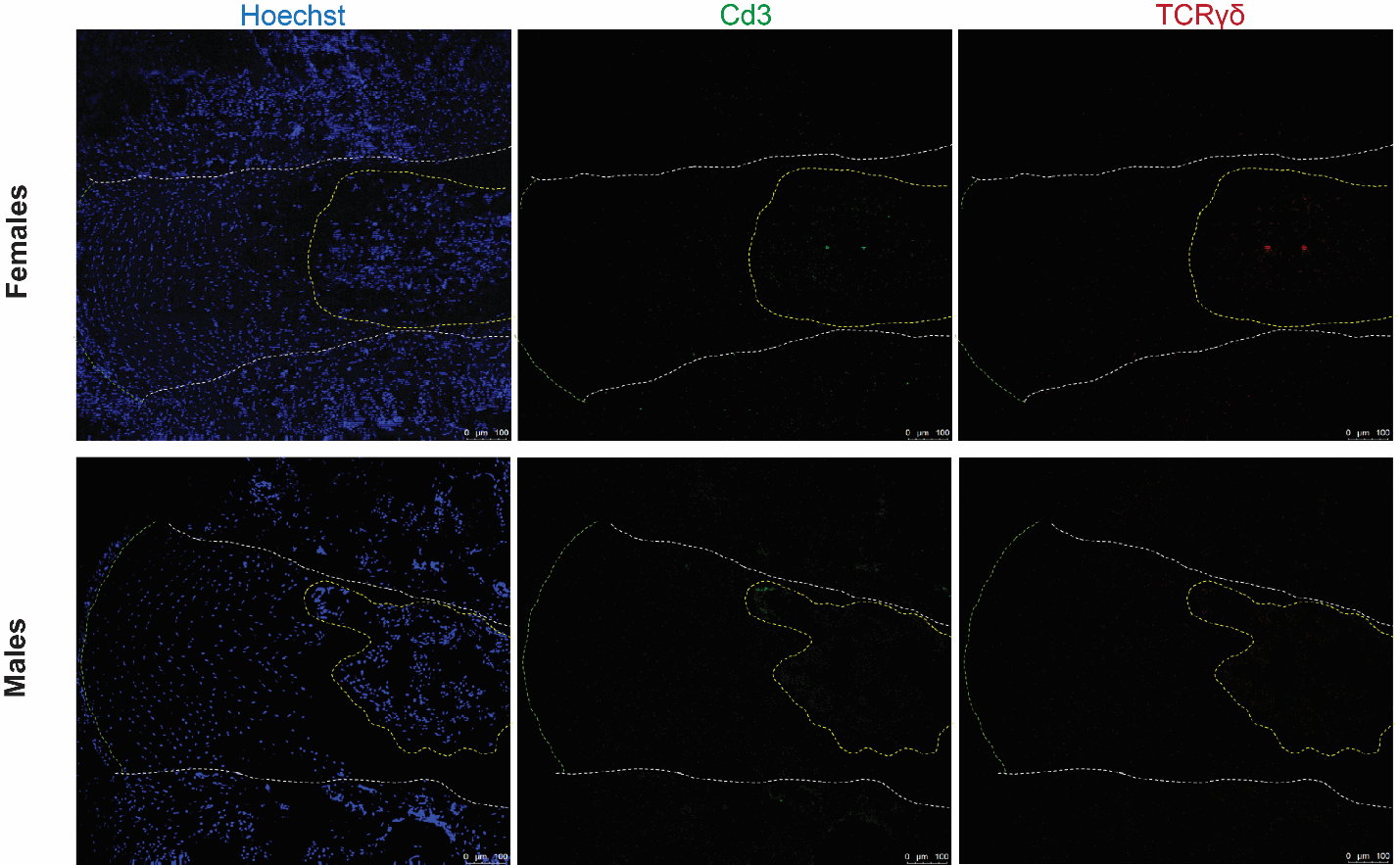

**Figure S5: Single channel analysis of Cd3 and TCRγδ immunofluorescence.** Separated immunofluorescence channels of Hoechst (nuclei), Cd3, and TCRγδ show the presence of γδ T cells in female but not male injured IVDs.

| **Table S1** | **Gene Name** | **Forward Primer: 5’-3’** | **Reverse Primer: 5’-3’** |
| --- | --- | --- | --- |
| 1 | Cd11b | CATCAAGGGCAGCCAGATTG | GGCCCCAATGAGGATCAAGT |
| 2 | Cd19 | GTCATTGCAAGGTCAGCAGTGTG | GAGAAAAGCCACCAGAGAAACC |
| 3 | Cd3g | TTCGCCAGTCAAGAGCTTCAG | CATATTCCCGGTCCTTGAGGG |
| 4 | Vegfa | CTTGTTCAGAGCGGAGAAAGC | ACATCTGCAAGTACGTTCGTT |
| 5 | Pdgfα | GATAGACTCCGTAGGGGCTGA | TCTCGGGCACATGGTTAATGG |
| 6 | Hprt | TCAGTCAACGGGGGACATAAA | GGGGCTGTACTGCTTAACCAG |
| 7 | Gapdh | AGAACATCATCCCTGCATCC | AGTTGCTGTTGAAGTCGC |

**Table S1: RTqPCR primers sequences**

| **Table S2:** | **Cd11b: Grouped: RM Two-way ANOVA (columns)** |
| --- | --- |

| Fixed effects (type III) | | P value | P value summary | | Statistically significant (P < 0.05)? | | F (DFn, DFd) | |
| --- | --- | --- | --- | --- | --- | --- | --- | --- |
| dpi | | 0.0044 | ** | | Yes | | F (10, 28) = 3.479 | |
| Injury | | <0.0001 | **** | | Yes | | F (1, 28) = 32.89 | |
| dpi x Injury | | 0.0044 | ** | | Yes | | F (10, 28) = 3.479 | |
| Holm-Šídák's multiple comparisons test | | Predicted (LS) mean diff. | Below threshold? | | Summary | | Adjusted P Value | |
| **Control - Stab** | |  |  | |  | |  | |
| 1 dpi | | -5.431 | Yes | | * | | 0.0409 | |
| 3 dpi | | -8.93 | Yes | | *** | | 0.0003 | |
| 5 dpi | | -3.516 | No | | ns | | 0.3701 | |
| 7 dpi | | -1.534 | No | | ns | | 0.9495 | |
| 10 dpi | | 0.3955 | No | | ns | | 0.9869 | |
| 12 dpi | | -10.18 | Yes | | *** | | 0.0003 | |
| 14 dpi | | -0.6174 | No | | ns | | 0.9869 | |
| 17 dpi | | -0.4517 | No | | ns | | 0.9869 | |
| 19 dpi | | -3.249 | No | | ns | | 0.5979 | |
| 21 dpi | | -1.318 | No | | ns | | 0.9753 | |
| 42 dpi | | -1.187 | No | | ns | | 0.9753 | |
| **Stab** | |  |  | |  | |  | |
| 1 dpi vs. 3 dpi | | -3.499 | No | | ns | | 0.8109 | |
| 1 dpi vs. 5 dpi | | 1.915 | No | | ns | | 0.9991 | |
| 1 dpi vs. 7 dpi | | 3.897 | No | | ns | | 0.6557 | |
| 1 dpi vs. 10 dpi | | 5.827 | No | | ns | | 0.0664 | |
| 1 dpi vs. 12 dpi | | -4.745 | No | | ns | | 0.4359 | |
| 1 dpi vs. 14 dpi | | 4.814 | No | | ns | | 0.4154 | |
| 1 dpi vs. 17 dpi | | 4.979 | No | | ns | | 0.2223 | |
| 1 dpi vs. 19 dpi | | 2.182 | No | | ns | | 0.9991 | |
| 1 dpi vs. 21 dpi | | 4.113 | No | | ns | | 0.6825 | |
| 1 dpi vs. 42 dpi | | 4.244 | No | | ns | | 0.6557 | |
| 3 dpi vs. 5 dpi | | 5.414 | No | | ns | | 0.1244 | |
| 3 dpi vs. 7 dpi | | 7.396 | Yes | | ** | | 0.0046 | |
| 3 dpi vs. 10 dpi | | 9.325 | Yes | | *** | | 0.0001 | |
| 3 dpi vs. 12 dpi | | -1.247 | No | | ns | | >0.9999 | |
| 3 dpi vs. 14 dpi | | 8.312 | Yes | | ** | | 0.0027 | |
| 3 dpi vs. 17 dpi | | 8.478 | Yes | | *** | | 0.0006 | |
| 3 dpi vs. 19 dpi | | 5.681 | No | | ns | | 0.1531 | |
| 3 dpi vs. 21 dpi | | 7.612 | Yes | | ** | | 0.0084 | |
| 3 dpi vs. 42 dpi | | 7.743 | Yes | | ** | | 0.0068 | |
| 5 dpi vs. 7 dpi | | 1.982 | No | | ns | | 0.9991 | |
| 5 dpi vs. 10 dpi | | 3.911 | No | | ns | | 0.6557 | |
| 5 dpi vs. 12 dpi | | -6.66 | Yes | | * | | 0.0395 | |
| 5 dpi vs. 14 dpi | | 2.899 | No | | ns | | 0.9822 | |
| 5 dpi vs. 17 dpi | | 3.064 | No | | ns | | 0.9308 | |
| 5 dpi vs. 19 dpi | | 0.267 | No | | ns | | >0.9999 | |
| 5 dpi vs. 21 dpi | | 2.198 | No | | ns | | 0.9991 | |
| 5 dpi vs. 42 dpi | | 2.329 | No | | ns | | 0.9984 | |
| 7 dpi vs. 10 dpi | | 1.929 | No | | ns | | 0.9991 | |
| 7 dpi vs. 12 dpi | | -8.643 | Yes | | ** | | 0.0015 | |
| 7 dpi vs. 14 dpi | | 0.9165 | No | | ns | | >0.9999 | |
| 7 dpi vs. 17 dpi | | 1.082 | No | | ns | | >0.9999 | |
| 7 dpi vs. 19 dpi | | -1.715 | No | | ns | | 0.9996 | |
| 7 dpi vs. 21 dpi | | 0.2161 | No | | ns | | >0.9999 | |
| 7 dpi vs. 42 dpi | | 0.3468 | No | | ns | | >0.9999 | |
| 10 dpi vs. 12 dpi | | -10.57 | Yes | | **** | | <0.0001 | |
| 10 dpi vs. 14 dpi | | -1.013 | No | | ns | | >0.9999 | |
| 10 dpi vs. 17 dpi | | -0.8472 | No | | ns | | >0.9999 | |
| 10 dpi vs. 19 dpi | | -3.644 | No | | ns | | 0.8485 | |
| 10 dpi vs. 21 dpi | | -1.713 | No | | ns | | 0.9996 | |
| 10 dpi vs. 42 dpi | | -1.582 | No | | ns | | 0.9996 | |
| 12 dpi vs. 14 dpi | | 9.559 | Yes | | *** | | 0.0009 | |
| 12 dpi vs. 17 dpi | | 9.725 | Yes | | *** | | 0.0002 | |
| 12 dpi vs. 19 dpi | | 6.927 | No | | ns | | 0.0511 | |
| 12 dpi vs. 21 dpi | | 8.859 | Yes | | ** | | 0.0028 | |
| 12 dpi vs. 42 dpi | | 8.989 | Yes | | ** | | 0.0023 | |
| 14 dpi vs. 17 dpi | | 0.1657 | No | | ns | | >0.9999 | |
| 14 dpi vs. 19 dpi | | -2.632 | No | | ns | | 0.9972 | |
| 14 dpi vs. 21 dpi | | -0.7004 | No | | ns | | >0.9999 | |
| 14 dpi vs. 42 dpi | | -0.5696 | No | | ns | | >0.9999 | |
| 17 dpi vs. 19 dpi | | -2.797 | No | | ns | | 0.9867 | |
| 17 dpi vs. 21 dpi | | -0.866 | No | | ns | | >0.9999 | |
| 17 dpi vs. 42 dpi | | -0.7353 | No | | ns | | >0.9999 | |
| 19 dpi vs. 21 dpi | | 1.931 | No | | ns | | 0.9995 | |
| 19 dpi vs. 42 dpi | | 2.062 | No | | ns | | 0.9993 | |
| 21 dpi vs. 42 dpi | | 0.1307 | No | | ns | | >0.9999 | |
| Table S2 | **Cd19: Grouped: RM Two-way ANOVA (columns)** | | | | | | | |
| Fixed effects (type III) | P value | | | P value summary | | Statistically significant (P < 0.05)? | | F (DFn, DFd) |
| dpi | <0.0001 | | | **** | | Yes | | F (10, 56) = 8.475 |
| Injury | 0.0002 | | | *** | | Yes | | F (1, 56) = 16.38 |
| dpi x Injury | <0.0001 | | | **** | | Yes | | F (10, 56) = 8.475 |
| Holm-Šídák's multiple comparisons test | Predicted (LS) mean diff. | | | Below threshold? | | Summary | | Adjusted P Value |
| **Control - Stab** |  | | |  | |  | |  |
| 1 dpi | -5.061 | | | Yes | | *** | | 0.0004 |
| 3 dpi | -10.13 | | | Yes | | **** | | <0.0001 |
| 5 dpi | 0.7309 | | | No | | ns | | 0.9945 |
| 7 dpi | -0.8328 | | | No | | ns | | 0.9936 |
| 10 dpi | 0.5089 | | | No | | ns | | 0.9945 |
| 12 dpi | 0.1207 | | | No | | ns | | 0.9945 |
| 14 dpi | -2.632 | | | No | | ns | | 0.3728 |
| 17 dpi | 0.7192 | | | No | | ns | | 0.9945 |
| 19 dpi | 0.6666 | | | No | | ns | | 0.9945 |
| 21 dpi | -0.33 | | | No | | ns | | 0.9945 |
| 42 dpi | -0.1828 | | | No | | ns | | 0.9945 |
| **Stab** |  | | |  | |  | |  |
| 1 dpi vs. 3 dpi | -5.074 | | | Yes | | ** | | 0.0017 |
| 1 dpi vs. 5 dpi | 5.792 | | | Yes | | *** | | 0.0002 |
| 1 dpi vs. 7 dpi | 4.228 | | | Yes | | * | | 0.0182 |
| 1 dpi vs. 10 dpi | 5.57 | | | Yes | | *** | | 0.0004 |
| 1 dpi vs. 12 dpi | 5.182 | | | Yes | | ** | | 0.0039 |
| 1 dpi vs. 14 dpi | 2.429 | | | No | | ns | | 0.8286 |
| 1 dpi vs. 17 dpi | 5.78 | | | Yes | | *** | | 0.0002 |
| 1 dpi vs. 19 dpi | 5.728 | | | Yes | | *** | | 0.0009 |
| 1 dpi vs. 21 dpi | 4.731 | | | Yes | | * | | 0.0122 |
| 1 dpi vs. 42 dpi | 4.878 | | | Yes | | ** | | 0.0085 |
| 3 dpi vs. 5 dpi | 10.87 | | | Yes | | **** | | <0.0001 |
| 3 dpi vs. 7 dpi | 9.302 | | | Yes | | **** | | <0.0001 |
| 3 dpi vs. 10 dpi | 10.64 | | | Yes | | **** | | <0.0001 |
| 3 dpi vs. 12 dpi | 10.26 | | | Yes | | **** | | <0.0001 |
| 3 dpi vs. 14 dpi | 7.502 | | | Yes | | **** | | <0.0001 |
| 3 dpi vs. 17 dpi | 10.85 | | | Yes | | **** | | <0.0001 |
| 3 dpi vs. 19 dpi | 10.8 | | | Yes | | **** | | <0.0001 |
| 3 dpi vs. 21 dpi | 9.805 | | | Yes | | **** | | <0.0001 |
| 3 dpi vs. 42 dpi | 9.952 | | | Yes | | **** | | <0.0001 |
| 5 dpi vs. 7 dpi | -1.564 | | | No | | ns | | 0.9956 |
| 5 dpi vs. 10 dpi | -0.2219 | | | No | | ns | | >0.9999 |
| 5 dpi vs. 12 dpi | -0.6101 | | | No | | ns | | >0.9999 |
| 5 dpi vs. 14 dpi | -3.363 | | | No | | ns | | 0.2696 |
| 5 dpi vs. 17 dpi | -0.01168 | | | No | | ns | | >0.9999 |
| 5 dpi vs. 19 dpi | -0.06427 | | | No | | ns | | >0.9999 |
| 5 dpi vs. 21 dpi | -1.061 | | | No | | ns | | >0.9999 |
| 5 dpi vs. 42 dpi | -0.9137 | | | No | | ns | | >0.9999 |
| 7 dpi vs. 10 dpi | 1.342 | | | No | | ns | | 0.9991 |
| 7 dpi vs. 12 dpi | 0.9535 | | | No | | ns | | >0.9999 |
| 7 dpi vs. 14 dpi | -1.799 | | | No | | ns | | 0.9909 |
| 7 dpi vs. 17 dpi | 1.552 | | | No | | ns | | 0.9956 |
| 7 dpi vs. 19 dpi | 1.499 | | | No | | ns | | 0.9988 |
| 7 dpi vs. 21 dpi | 0.5028 | | | No | | ns | | >0.9999 |
| 7 dpi vs. 42 dpi | 0.65 | | | No | | ns | | >0.9999 |
| 10 dpi vs. 12 dpi | -0.3882 | | | No | | ns | | >0.9999 |
| 10 dpi vs. 14 dpi | -3.141 | | | No | | ns | | 0.3796 |
| 10 dpi vs. 17 dpi | 0.2102 | | | No | | ns | | >0.9999 |
| 10 dpi vs. 19 dpi | 0.1577 | | | No | | ns | | >0.9999 |
| 10 dpi vs. 21 dpi | -0.8389 | | | No | | ns | | >0.9999 |
| 10 dpi vs. 42 dpi | -0.6917 | | | No | | ns | | >0.9999 |
| 12 dpi vs. 14 dpi | -2.753 | | | No | | ns | | 0.7501 |
| 12 dpi vs. 17 dpi | 0.5985 | | | No | | ns | | >0.9999 |
| 12 dpi vs. 19 dpi | 0.5459 | | | No | | ns | | >0.9999 |
| 12 dpi vs. 21 dpi | -0.4507 | | | No | | ns | | >0.9999 |
| 12 dpi vs. 42 dpi | -0.3035 | | | No | | ns | | >0.9999 |
| 14 dpi vs. 17 dpi | 3.351 | | | No | | ns | | 0.2696 |
| 14 dpi vs. 19 dpi | 3.299 | | | No | | ns | | 0.4058 |
| 14 dpi vs. 21 dpi | 2.302 | | | No | | ns | | 0.9325 |
| 14 dpi vs. 42 dpi | 2.449 | | | No | | ns | | 0.8879 |
| 17 dpi vs. 19 dpi | -0.05258 | | | No | | ns | | >0.9999 |
| 17 dpi vs. 21 dpi | -1.049 | | | No | | ns | | >0.9999 |
| 17 dpi vs. 42 dpi | -0.902 | | | No | | ns | | >0.9999 |
| 19 dpi vs. 21 dpi | -0.9966 | | | No | | ns | | >0.9999 |
| 19 dpi vs. 42 dpi | -0.8494 | | | No | | ns | | >0.9999 |
| 21 dpi vs. 42 dpi | 0.1472 | | | No | | ns | | >0.9999 |

| Tables S2 | **Cd3g: Grouped: RM Two-way ANOVA (columns)** |
| --- | --- |

| Fixed effects (type III) | P value | | | P value summary | | Statistically significant (P < 0.05)? | | | F (DFn, DFd) | |
| --- | --- | --- | --- | --- | --- | --- | --- | --- | --- | --- |
| dpi | <0.0001 | | | **** | | Yes | | | F (10, 50) = 11.55 | |
| injury | <0.0001 | | | **** | | Yes | | | F (1, 50) = 58.13 | |
| dpi x injury | <0.0001 | | | **** | | Yes | | | F (10, 50) = 11.55 | |
| Holm-Šídák's multiple comparisons test | Predicted (LS) mean diff. | | | Below threshold? | | Summary | | | Adjusted P Value | |
| **Control - Stab** |  | | |  | |  | | |  | |
| 1 dpi | -3.205 | | | No | | ns | | | 0.958 | |
| 3 dpi | -14.97 | | | Yes | | ** | | | 0.0019 | |
| 5 dpi | -0.5843 | | | No | | ns | | | 0.997 | |
| 7 dpi | -1.745 | | | No | | ns | | | 0.9954 | |
| 10 dpi | -0.3035 | | | No | | ns | | | 0.997 | |
| 12 dpi | -1.89 | | | No | | ns | | | 0.9954 | |
| 14 dpi | -3.225 | | | No | | ns | | | 0.9696 | |
| 17 dpi | -0.5919 | | | No | | ns | | | 0.997 | |
| 19 dpi | -26.32 | | | Yes | | **** | | | <0.0001 | |
| 21 dpi | -37.25 | | | Yes | | **** | | | <0.0001 | |
| 42 dpi | -1.223 | | | No | | ns | | | 0.9958 | |
| **Stab** |  | | |  | |  | | |  | |
| 1 dpi vs. 3 dpi | -11.76 | | | Yes | | * | | | 0.0466 | |
| 1 dpi vs. 5 dpi | 2.621 | | | No | | ns | | | >0.9999 | |
| 1 dpi vs. 7 dpi | 1.46 | | | No | | ns | | | >0.9999 | |
| 1 dpi vs. 10 dpi | 2.901 | | | No | | ns | | | >0.9999 | |
| 1 dpi vs. 12 dpi | 1.315 | | | No | | ns | | | >0.9999 | |
| 1 dpi vs. 14 dpi | -0.01976 | | | No | | ns | | | >0.9999 | |
| 1 dpi vs. 17 dpi | 2.613 | | | No | | ns | | | >0.9999 | |
| 1 dpi vs. 19 dpi | -23.11 | | | Yes | | **** | | | <0.0001 | |
| 1 dpi vs. 21 dpi | -34.05 | | | Yes | | **** | | | <0.0001 | |
| 1 dpi vs. 42 dpi | 1.982 | | | No | | ns | | | >0.9999 | |
| 3 dpi vs. 5 dpi | 14.38 | | | Yes | | * | | | 0.0122 | |
| 3 dpi vs. 7 dpi | 13.22 | | | Yes | | * | | | 0.0146 | |
| 3 dpi vs. 10 dpi | 14.66 | | | Yes | | ** | | | 0.0099 | |
| 3 dpi vs. 12 dpi | 13.08 | | | Yes | | * | | | 0.0325 | |
| 3 dpi vs. 14 dpi | 11.74 | | | No | | ns | | | 0.0844 | |
| 3 dpi vs. 17 dpi | 14.38 | | | Yes | | ** | | | 0.0056 | |
| 3 dpi vs. 19 dpi | -11.35 | | | No | | ns | | | 0.1086 | |
| 3 dpi vs. 21 dpi | -22.29 | | | Yes | | **** | | | <0.0001 | |
| 3 dpi vs. 42 dpi | 13.75 | | | Yes | | * | | | 0.0196 | |
| 5 dpi vs. 7 dpi | -1.16 | | | No | | ns | | | >0.9999 | |
| 5 dpi vs. 10 dpi | 0.2808 | | | No | | ns | | | >0.9999 | |
| 5 dpi vs. 12 dpi | -1.306 | | | No | | ns | | | >0.9999 | |
| 5 dpi vs. 14 dpi | -2.64 | | | No | | ns | | | >0.9999 | |
| 5 dpi vs. 17 dpi | -0.00755 | | | No | | ns | | | >0.9999 | |
| 5 dpi vs. 19 dpi | -25.73 | | | Yes | | **** | | | <0.0001 | |
| 5 dpi vs. 21 dpi | -36.67 | | | Yes | | **** | | | <0.0001 | |
| 5 dpi vs. 42 dpi | -0.6383 | | | No | | ns | | | >0.9999 | |
| 7 dpi vs. 10 dpi | 1.441 | | | No | | ns | | | >0.9999 | |
| 7 dpi vs. 12 dpi | -0.1453 | | | No | | ns | | | >0.9999 | |
| 7 dpi vs. 14 dpi | -1.48 | | | No | | ns | | | >0.9999 | |
| 7 dpi vs. 17 dpi | 1.153 | | | No | | ns | | | >0.9999 | |
| 7 dpi vs. 19 dpi | -24.57 | | | Yes | | **** | | | <0.0001 | |
| 7 dpi vs. 21 dpi | -35.51 | | | Yes | | **** | | | <0.0001 | |
| 7 dpi vs. 42 dpi | 0.5221 | | | No | | ns | | | >0.9999 | |
| 10 dpi vs. 12 dpi | -1.587 | | | No | | ns | | | >0.9999 | |
| 10 dpi vs. 14 dpi | -2.921 | | | No | | ns | | | >0.9999 | |
| 10 dpi vs. 17 dpi | -0.2883 | | | No | | ns | | | >0.9999 | |
| 10 dpi vs. 19 dpi | -26.01 | | | Yes | | **** | | | <0.0001 | |
| 10 dpi vs. 21 dpi | -36.95 | | | Yes | | **** | | | <0.0001 | |
| 10 dpi vs. 42 dpi | -0.9191 | | | No | | ns | | | >0.9999 | |
| 12 dpi vs. 14 dpi | -1.335 | | | No | | ns | | | >0.9999 | |
| 12 dpi vs. 17 dpi | 1.298 | | | No | | ns | | | >0.9999 | |
| 12 dpi vs. 19 dpi | -24.43 | | | Yes | | **** | | | <0.0001 | |
| 12 dpi vs. 21 dpi | -35.36 | | | Yes | | **** | | | <0.0001 | |
| 12 dpi vs. 42 dpi | 0.6674 | | | No | | ns | | | >0.9999 | |
| 14 dpi vs. 17 dpi | 2.633 | | | No | | ns | | | >0.9999 | |
| 14 dpi vs. 19 dpi | -23.09 | | | Yes | | **** | | | <0.0001 | |
| 14 dpi vs. 21 dpi | -34.03 | | | Yes | | **** | | | <0.0001 | |
| 14 dpi vs. 42 dpi | 2.002 | | | No | | ns | | | >0.9999 | |
| 17 dpi vs. 19 dpi | -25.72 | | | Yes | | **** | | | <0.0001 | |
| 17 dpi vs. 21 dpi | -36.66 | | | Yes | | **** | | | <0.0001 | |
| 17 dpi vs. 42 dpi | -0.6307 | | | No | | ns | | | >0.9999 | |
| 19 dpi vs. 21 dpi | -10.94 | | | No | | ns | | | 0.1398 | |
| 19 dpi vs. 42 dpi | 25.09 | | | Yes | | **** | | | <0.0001 | |
| 21 dpi vs. 42 dpi | 36.03 | | | Yes | | **** | | | <0.0001 | |
| Table S2 | **Vegfa: Grouped: RM Two-way ANOVA (columns)** | | | | | | | | | |
| Fixed effects (type III) | P value | | P value summary | | Statistically significant (P < 0.05)? | | | F (DFn, DFd) | | |
| dpi | <0.0001 | | **** | | Yes | | | F (10, 56) = 11.08 | | |
| injury | <0.0001 | | **** | | Yes | | | F (1, 56) = 31.28 | | |
| dpi x injury | <0.0001 | | **** | | Yes | | | F (10, 56) = 11.08 | | |
| Holm-Šídák's multiple comparisons test | Predicted (LS) mean diff. | | Below threshold? | | Summary | | | Adjusted P Value | | |
| **Control - Stab** |  | |  | |  | | |  | | |
| 1 dpi | 0.1704 | | No | | ns | | | 0.9987 | | |
| 3 dpi | 0.6219 | | No | | ns | | | 0.9987 | | |
| 5 dpi | 0.7567 | | No | | ns | | | 0.9985 | | |
| 7 dpi | 0.5638 | | No | | ns | | | 0.9987 | | |
| 10 dpi | -3.955 | | Yes | | * | | | 0.0351 | | |
| 12 dpi | 0.4434 | | No | | ns | | | 0.9987 | | |
| 14 dpi | 0.2851 | | No | | ns | | | 0.9987 | | |
| 17 dpi | -21.28 | | Yes | | **** | | | <0.0001 | | |
| 19 dpi | -4.675 | | No | | ns | | | 0.0635 | | |
| 21 dpi | -0.1142 | | No | | ns | | | 0.9987 | | |
| 42 dpi | -2.416 | | No | | ns | | | 0.7414 | | |
| **Stab** |  | |  | |  | | |  | | |
| 1 dpi vs. 3 dpi | 0.4515 | | No | | ns | | | >0.9999 | | |
| 1 dpi vs. 5 dpi | 0.5863 | | No | | ns | | | >0.9999 | | |
| 1 dpi vs. 7 dpi | 0.3934 | | No | | ns | | | >0.9999 | | |
| 1 dpi vs. 10 dpi | -4.125 | | No | | ns | | | 0.0887 | | |
| 1 dpi vs. 12 dpi | 0.273 | | No | | ns | | | >0.9999 | | |
| 1 dpi vs. 14 dpi | 0.1147 | | No | | ns | | | >0.9999 | | |
| 1 dpi vs. 17 dpi | -21.45 | | Yes | | **** | | | <0.0001 | | |
| 1 dpi vs. 19 dpi | -4.846 | | No | | ns | | | 0.078 | | |
| 1 dpi vs. 21 dpi | -0.2846 | | No | | ns | | | >0.9999 | | |
| 1 dpi vs. 42 dpi | -2.586 | | No | | ns | | | 0.9295 | | |
| 3 dpi vs. 5 dpi | 0.1348 | | No | | ns | | | >0.9999 | | |
| 3 dpi vs. 7 dpi | -0.05813 | | No | | ns | | | >0.9999 | | |
| 3 dpi vs. 10 dpi | -4.577 | | No | | ns | | | 0.063 | | |
| 3 dpi vs. 12 dpi | -0.1785 | | No | | ns | | | >0.9999 | | |
| 3 dpi vs. 14 dpi | -0.3367 | | No | | ns | | | >0.9999 | | |
| 3 dpi vs. 17 dpi | -21.91 | | Yes | | **** | | | <0.0001 | | |
| 3 dpi vs. 19 dpi | -5.297 | | No | | ns | | | 0.0563 | | |
| 3 dpi vs. 21 dpi | -0.736 | | No | | ns | | | >0.9999 | | |
| 3 dpi vs. 42 dpi | -3.037 | | No | | ns | | | 0.8335 | | |
| 5 dpi vs. 7 dpi | -0.1929 | | No | | ns | | | >0.9999 | | |
| 5 dpi vs. 10 dpi | -4.712 | | No | | ns | | | 0.0507 | | |
| 5 dpi vs. 12 dpi | -0.3133 | | No | | ns | | | >0.9999 | | |
| 5 dpi vs. 14 dpi | -0.4715 | | No | | ns | | | >0.9999 | | |
| 5 dpi vs. 17 dpi | -22.04 | | Yes | | **** | | | <0.0001 | | |
| 5 dpi vs. 19 dpi | -5.432 | | Yes | | * | | | 0.0456 | | |
| 5 dpi vs. 21 dpi | -0.8708 | | No | | ns | | | >0.9999 | | |
| 5 dpi vs. 42 dpi | -3.172 | | No | | ns | | | 0.7827 | | |
| 7 dpi vs. 10 dpi | -4.519 | | No | | ns | | | 0.0695 | | |
| 7 dpi vs. 12 dpi | -0.1204 | | No | | ns | | | >0.9999 | | |
| 7 dpi vs. 14 dpi | -0.2786 | | No | | ns | | | >0.9999 | | |
| 7 dpi vs. 17 dpi | -21.85 | | Yes | | **** | | | <0.0001 | | |
| 7 dpi vs. 19 dpi | -5.239 | | No | | ns | | | 0.0613 | | |
| 7 dpi vs. 21 dpi | -0.6779 | | No | | ns | | | >0.9999 | | |
| 7 dpi vs. 42 dpi | -2.979 | | No | | ns | | | 0.8476 | | |
| 10 dpi vs. 12 dpi | 4.398 | | No | | ns | | | 0.1575 | | |
| 10 dpi vs. 14 dpi | 4.24 | | No | | ns | | | 0.1997 | | |
| 10 dpi vs. 17 dpi | -17.33 | | Yes | | **** | | | <0.0001 | | |
| 10 dpi vs. 19 dpi | -0.7201 | | No | | ns | | | >0.9999 | | |
| 10 dpi vs. 21 dpi | 3.841 | | No | | ns | | | 0.3467 | | |
| 10 dpi vs. 42 dpi | 1.539 | | No | | ns | | | 0.9998 | | |
| 12 dpi vs. 14 dpi | -0.1582 | | No | | ns | | | >0.9999 | | |
| 12 dpi vs. 17 dpi | -21.73 | | Yes | | **** | | | <0.0001 | | |
| 12 dpi vs. 19 dpi | -5.119 | | No | | ns | | | 0.1212 | | |
| 12 dpi vs. 21 dpi | -0.5575 | | No | | ns | | | >0.9999 | | |
| 12 dpi vs. 42 dpi | -2.859 | | No | | ns | | | 0.9303 | | |
| 14 dpi vs. 17 dpi | -21.57 | | Yes | | **** | | | <0.0001 | | |
| 14 dpi vs. 19 dpi | -4.96 | | No | | ns | | | 0.1514 | | |
| 14 dpi vs. 21 dpi | -0.3993 | | No | | ns | | | >0.9999 | | |
| 14 dpi vs. 42 dpi | -2.701 | | No | | ns | | | 0.9559 | | |
| 17 dpi vs. 19 dpi | 16.61 | | Yes | | **** | | | <0.0001 | | |
| 17 dpi vs. 21 dpi | 21.17 | | Yes | | **** | | | <0.0001 | | |
| 17 dpi vs. 42 dpi | 18.87 | | Yes | | **** | | | <0.0001 | | |
| 19 dpi vs. 21 dpi | 4.561 | | No | | ns | | | 0.2509 | | |
| 19 dpi vs. 42 dpi | 2.26 | | No | | ns | | | 0.9923 | | |
| 21 dpi vs. 42 dpi | -2.301 | | No | | ns | | | 0.992 | | |
| Table S2 | **Pdgfa: Grouped: RM Two-way ANOVA (columns)** | | | | | | | | | |
| Fixed effects (type III) | P value | P value summary | | | | | Statistically significant (P < 0.05)? | | | F (DFn, DFd) |
| dpi | 0.0058 | ** | | | | | Yes | | | F (10, 44) = 2.992 |
| injury | <0.0001 | **** | | | | | Yes | | | F (1, 44) = 22.98 |
| dpi x injury | 0.0058 | ** | | | | | Yes | | | F (10, 44) = 2.992 |
| Holm-Šídák's multiple comparisons test | Predicted (LS) mean diff. | Below threshold? | | | | | Summary | | | Adjusted P Value |
| **Control - Stab** |  |  | | | | |  | | |  |
| 1 dpi | 0.9272 | No | | | | | ns | | | 0.9605 |
| 3 dpi | -0.263 | No | | | | | ns | | | 0.9842 |
| 5 dpi | 0.4148 | No | | | | | ns | | | 0.9842 |
| 7 dpi | -2.482 | No | | | | | ns | | | 0.3134 |
| 10 dpi | 0.652 | No | | | | | ns | | | 0.9782 |
| 12 dpi | -2.887 | No | | | | | ns | | | 0.1935 |
| 14 dpi | -4.623 | Yes | | | | | ** | | | 0.0084 |
| 17 dpi | -3.548 | No | | | | | ns | | | 0.0747 |
| 19 dpi | -5.286 | Yes | | | | | ** | | | 0.0019 |
| 21 dpi | -3.158 | No | | | | | ns | | | 0.1378 |
| 42 dpi | -0.2482 | No | | | | | ns | | | 0.9842 |
| **Stab** |  |  | | | | |  | | |  |
| 1 dpi vs. 3 dpi | -1.19 | -5.565 to 3.185 | | | | | No | | | ns |
| 1 dpi vs. 5 dpi | -0.5124 | -4.887 to 3.862 | | | | | No | | | ns |
| 1 dpi vs. 7 dpi | -3.409 | -7.784 to 0.9660 | | | | | No | | | ns |
| 1 dpi vs. 10 dpi | -0.2751 | -4.650 to 4.100 | | | | | No | | | ns |
| 1 dpi vs. 12 dpi | -3.814 | -8.189 to 0.5606 | | | | | No | | | ns |
| 1 dpi vs. 14 dpi | -5.55 | -9.925 to -1.175 | | | | | Yes | | | ** |
| 1 dpi vs. 17 dpi | -4.475 | -8.850 to -0.09981 | | | | | Yes | | | * |
| 1 dpi vs. 19 dpi | -6.213 | -10.59 to -1.838 | | | | | Yes | | | *** |
| 1 dpi vs. 21 dpi | -4.085 | -8.460 to 0.2897 | | | | | No | | | ns |
| 1 dpi vs. 42 dpi | -1.175 | -5.550 to 3.199 | | | | | No | | | ns |
| 3 dpi vs. 5 dpi | 0.6778 | -3.697 to 5.053 | | | | | No | | | ns |
| 3 dpi vs. 7 dpi | -2.219 | -6.594 to 2.156 | | | | | No | | | ns |
| 3 dpi vs. 10 dpi | 0.915 | -3.460 to 5.290 | | | | | No | | | ns |
| 3 dpi vs. 12 dpi | -2.624 | -6.999 to 1.751 | | | | | No | | | ns |
| 3 dpi vs. 14 dpi | -4.36 | -8.735 to 0.01494 | | | | | No | | | ns |
| 3 dpi vs. 17 dpi | -3.285 | -7.659 to 1.090 | | | | | No | | | ns |
| 3 dpi vs. 19 dpi | -5.023 | -9.398 to -0.6481 | | | | | Yes | | | * |
| 3 dpi vs. 21 dpi | -2.895 | -7.270 to 1.480 | | | | | No | | | ns |
| 3 dpi vs. 42 dpi | 0.01472 | -4.360 to 4.390 | | | | | No | | | ns |
| 5 dpi vs. 7 dpi | -2.896 | -7.271 to 1.478 | | | | | No | | | ns |
| 5 dpi vs. 10 dpi | 0.2372 | -4.138 to 4.612 | | | | | No | | | ns |
| 5 dpi vs. 12 dpi | -3.302 | -7.677 to 1.073 | | | | | No | | | ns |
| 5 dpi vs. 14 dpi | -5.038 | -9.413 to -0.6628 | | | | | Yes | | | * |
| 5 dpi vs. 17 dpi | -3.962 | -8.337 to 0.4125 | | | | | No | | | ns |
| 5 dpi vs. 19 dpi | -5.701 | -10.08 to -1.326 | | | | | Yes | | | ** |
| 5 dpi vs. 21 dpi | -3.573 | -7.948 to 0.8021 | | | | | No | | | ns |
| 5 dpi vs. 42 dpi | -0.663 | -5.038 to 3.712 | | | | | No | | | ns |
| 7 dpi vs. 10 dpi | 3.134 | -1.241 to 7.509 | | | | | No | | | ns |
| 7 dpi vs. 12 dpi | -0.4055 | -4.780 to 3.969 | | | | | No | | | ns |
| 7 dpi vs. 14 dpi | -2.141 | -6.516 to 2.234 | | | | | No | | | ns |
| 7 dpi vs. 17 dpi | -1.066 | -5.441 to 3.309 | | | | | No | | | ns |
| 7 dpi vs. 19 dpi | -2.804 | -7.179 to 1.571 | | | | | No | | | ns |
| 7 dpi vs. 21 dpi | -0.6763 | -5.051 to 3.699 | | | | | No | | | ns |
| 7 dpi vs. 42 dpi | 2.233 | -2.141 to 6.608 | | | | | No | | | ns |
| 10 dpi vs. 12 dpi | -3.539 | -7.914 to 0.8357 | | | | | No | | | ns |
| 10 dpi vs. 14 dpi | -5.275 | -9.650 to -0.9001 | | | | | Yes | | | ** |
| 10 dpi vs. 17 dpi | -4.2 | -8.574 to 0.1753 | | | | | No | | | ns |
| 10 dpi vs. 19 dpi | -5.938 | -10.31 to -1.563 | | | | | Yes | | | ** |
| 10 dpi vs. 21 dpi | -3.81 | -8.185 to 0.5649 | | | | | No | | | ns |
| 10 dpi vs. 42 dpi | -0.9003 | -5.275 to 3.475 | | | | | No | | | ns |
| 12 dpi vs. 14 dpi | -1.736 | -6.111 to 2.639 | | | | | No | | | ns |
| 12 dpi vs. 17 dpi | -0.6604 | -5.035 to 3.714 | | | | | No | | | ns |
| 12 dpi vs. 19 dpi | -2.399 | -6.774 to 1.976 | | | | | No | | | ns |
| 12 dpi vs. 21 dpi | -0.2708 | -4.646 to 4.104 | | | | | No | | | ns |
| 12 dpi vs. 42 dpi | 2.639 | -1.736 to 7.014 | | | | | No | | | ns |
| 14 dpi vs. 17 dpi | 1.075 | -3.299 to 5.450 | | | | | No | | | ns |
| 14 dpi vs. 19 dpi | -0.6631 | -5.038 to 3.712 | | | | | No | | | ns |
| 14 dpi vs. 21 dpi | 1.465 | -2.910 to 5.840 | | | | | No | | | ns |
| 14 dpi vs. 42 dpi | 4.375 | -0.0002202 to 8.749 | | | | | No | | | ns |
| 17 dpi vs. 19 dpi | -1.738 | -6.113 to 2.636 | | | | | No | | | ns |
| 17 dpi vs. 21 dpi | 0.3896 | -3.985 to 4.764 | | | | | No | | | ns |
| 17 dpi vs. 42 dpi | 3.299 | -1.076 to 7.674 | | | | | No | | | ns |
| 19 dpi vs. 21 dpi | 2.128 | -2.247 to 6.503 | | | | | No | | | ns |
| 19 dpi vs. 42 dpi | 5.038 | 0.6628 to 9.413 | | | | | Yes | | | * |
| 21 dpi vs. 42 dpi | 2.91 | -1.465 to 7.285 | | | | | No | | | ns |

**Table S2:** **Mixed design two way ANOVA fixed effects and post hoc results for female longitudinal data.**

| Table S3 | **Cd11b: Grouped: RM Two-way ANOVA (columns)** |
| --- | --- |

| Fixed effects (type III) | P value | P value summary | Statistically significant (P < 0.05)? | F (DFn, DFd) |
| --- | --- | --- | --- | --- |
| dpi | 0.0483 | * | Yes | F (4, 10) = 3.524 |
| Injury | 0.0005 | *** | Yes | F (1, 10) = 25.87 |
| dpi x Injury | 0.0483 | * | Yes | F (4, 10) = 3.524 |
| Holm-Šídák's multiple comparisons test | Predicted (LS) mean diff. | Below threshold? | Summary | Adjusted P Value |
| **Control - Stab** |  |  |  |  |
| 3 dpi | -3.062 | No | ns | 0.5099 |
| 7 dpi | -14.94 | Yes | ** | 0.0012 |
| 12 dpi | -5.545 | No | ns | 0.2375 |
| 19 dpi | -3.38 | No | ns | 0.5099 |
| 42 dpi | -3.58 | No | ns | 0.5099 |
| **Stab** |  |  |  |  |
| 3 dpi vs. 7 dpi | -11.88 | -19.90 to -3.850 | Yes | ** |
| 3 dpi vs. 12 dpi | -2.484 | -10.51 to 5.542 | No | ns |
| 3 dpi vs. 19 dpi | -0.3184 | -8.344 to 7.707 | No | ns |
| 3 dpi vs. 42 dpi | -0.518 | -8.544 to 7.508 | No | ns |
| 7 dpi vs. 12 dpi | 9.392 | 1.367 to 17.42 | Yes | * |
| 7 dpi vs. 19 dpi | 11.56 | 3.532 to 19.58 | Yes | ** |
| 7 dpi vs. 42 dpi | 11.36 | 3.332 to 19.38 | Yes | ** |
| 12 dpi vs. 19 dpi | 2.165 | -5.860 to 10.19 | No | ns |
| 12 dpi vs. 42 dpi | 1.966 | -6.060 to 9.991 | No | ns |
| 19 dpi vs. 42 dpi | -0.1997 | -8.225 to 7.826 | No | ns |

| Table S3 | **Cd19: Grouped: RM Two-way ANOVA (columns)** |
| --- | --- |

| Fixed effects (type III) | | P value | | P value summary | Statistically significant (P < 0.05)? | | F (DFn, DFd) |
| --- | --- | --- | --- | --- | --- | --- | --- |
| dpi | | 0.4396 | | ns | No | | F (4, 10) = 1.027 |
| Injury | | 0.8038 | | ns | No | | F (1, 10) = 0.06509 |
| dpi x Injury | | 0.4396 | | ns | No | | F (4, 10) = 1.027 |
| Holm-Šídák's multiple comparisons test | | Predicted (LS) mean diff. | | Below threshold? | Summary | | Adjusted P Value |
| **Control - Stab** | |  | |  |  | |  |
| 3 dpi | | -0.4754 | | No | ns | | 0.8913 |
| 7 dpi | | 0.1175 | | No | ns | | 0.8913 |
| 12 dpi | | 0.5732 | | No | ns | | 0.8913 |
| 19 dpi | | 0.4812 | | No | ns | | 0.8913 |
| 42 dpi | | -1.09 | | No | ns | | 0.5441 |
| Table S3 | | **Cd3g: Grouped: RM Two-way ANOVA (columns)** | | | | | |
| Fixed effects (type III) | P value | | P value summary | | Statistically significant (P < 0.05)? | F (DFn, DFd) | |
| dpi | 0.2945 | | ns | | No | F (4, 20) = 1.326 | |
| injury | 0.0003 | | *** | | Yes | F (1, 20) = 18.92 | |
| dpi x injury | 0.2945 | | ns | | No | F (4, 20) = 1.326 | |
| Holm-Šídák's multiple comparisons test | Predicted (LS) mean diff. | | Below threshold? | | Summary | Adjusted P Value | |
| **Control - Stab** |  | |  | |  |  | |
| 3 dpi | -3.427 | | -6.649 to -0.2051 | | Yes | * | |
| 7 dpi | -1.435 | | -4.657 to 1.786 | | No | ns | |
| 12 dpi | -0.52 | | -3.742 to 2.702 | | No | ns | |
| 19 dpi | -2.082 | | -5.304 to 1.140 | | No | ns | |
| 42 dpi | -3.586 | | -6.808 to -0.3641 | | Yes | * | |
| **Stab** |  | |  | |  |  | |
| 3 dpi vs. 7 dpi | 1.992 | | -1.408 to 5.391 | | No | ns | |
| 3 dpi vs. 12 dpi | 2.907 | | -0.4924 to 6.306 | | No | ns | |
| 3 dpi vs. 19 dpi | 1.345 | | -2.054 to 4.744 | | No | ns | |
| 3 dpi vs. 42 dpi | -0.159 | | -3.558 to 3.240 | | No | ns | |
| 7 dpi vs. 12 dpi | 0.9155 | | -2.484 to 4.315 | | No | ns | |
| 7 dpi vs. 19 dpi | -0.6465 | | -4.046 to 2.753 | | No | ns | |
| 7 dpi vs. 42 dpi | -2.151 | | -5.550 to 1.249 | | No | ns | |
| 12 dpi vs. 19 dpi | -1.562 | | -4.961 to 1.837 | | No | ns | |
| 12 dpi vs. 42 dpi | -3.066 | | -6.465 to 0.3334 | | No | ns | |
| 19 dpi vs. 42 dpi | -1.504 | | -4.903 to 1.895 | | No | ns | |

| Table S3 | **Vegfa: Grouped: RM Two-way ANOVA (columns)** |
| --- | --- |

| Fixed effects (type III) | P value | P value summary | Statistically significant (P < 0.05)? | F (DFn, DFd) |
| --- | --- | --- | --- | --- |
| dpi | 0.7908 | ns | No | F (4, 10) = 0.4201 |
| injury | <0.0001 | **** | Yes | F (1, 10) = 42.71 |
| dpi x injury | 0.7908 | ns | No | F (4, 10) = 0.4201 |
| Holm-Šídák's multiple comparisons test | Predicted (LS) mean diff. | Below threshold? | Summary | Adjusted P Value |
| **Control - Stab** |  |  |  |  |
| 3 dpi | 0.5745 | Yes | * | 0.0426 |
| 7 dpi | 0.5786 | Yes | * | 0.0426 |
| 12 dpi | 0.5762 | Yes | * | 0.0426 |
| 19 dpi | 0.5555 | Yes | * | 0.0426 |
| 42 dpi | 0.3142 | No | ns | 0.1077 |
| **Stab** |  |  |  |  |
| 3 dpi vs. 7 dpi | 0.004092 | -0.5281 to 0.5363 | No | ns |
| 3 dpi vs. 12 dpi | 0.001724 | -0.5305 to 0.5340 | No | ns |
| 3 dpi vs. 19 dpi | -0.01905 | -0.5513 to 0.5132 | No | ns |
| 3 dpi vs. 42 dpi | -0.2603 | -0.7925 to 0.2720 | No | ns |
| 7 dpi vs. 12 dpi | -0.00237 | -0.5346 to 0.5299 | No | ns |
| 7 dpi vs. 19 dpi | -0.02314 | -0.5554 to 0.5091 | No | ns |
| 7 dpi vs. 42 dpi | -0.2644 | -0.7966 to 0.2679 | No | ns |
| 12 dpi vs. 19 dpi | -0.02077 | -0.5530 to 0.5115 | No | ns |
| 12 dpi vs. 42 dpi | -0.262 | -0.7942 to 0.2702 | No | ns |
| 19 dpi vs. 42 dpi | -0.2412 | -0.7735 to 0.2910 | No | ns |

| Tables S3 | **Pdgfa: Grouped: RM Two-way ANOVA (columns)** |
| --- | --- |

| Fixed effects (type III) | P value | P value summary | Statistically significant (P < 0.05)? | F (DFn, DFd) |
| --- | --- | --- | --- | --- |
| dpi | 0.0215 | * | Yes | F (4, 20) = 3.661 |
| injury | 0.0492 | * | Yes | F (1, 20) = 4.383 |
| dpi x injury | 0.0215 | * | Yes | F (4, 20) = 3.661 |
| Holm-Šídák's multiple comparisons test | Predicted (LS) mean diff. | Below threshold? | Summary | Adjusted P Value |
| **Control - Stab** |  |  |  |  |
| 3 dpi | 0.2888 | No | ns | 0.7518 |
| 7 dpi | -1.063 | Yes | ** | 0.0038 |
| 12 dpi | -0.2836 | No | ns | 0.7518 |
| 19 dpi | 0.06703 | No | ns | 0.8051 |
| 42 dpi | -0.2647 | No | ns | 0.7518 |
| **Stab** |  |  |  |  |
| 3 dpi vs. 7 dpi | -1.352 | -2.154 to -0.5493 | Yes | *** |
| 3 dpi vs. 12 dpi | -0.5724 | -1.375 to 0.2299 | No | ns |
| 3 dpi vs. 19 dpi | -0.2218 | -1.024 to 0.5805 | No | ns |
| 3 dpi vs. 42 dpi | -0.5535 | -1.356 to 0.2488 | No | ns |
| 7 dpi vs. 12 dpi | 0.7791 | -0.02313 to 1.581 | No | ns |
| 7 dpi vs. 19 dpi | 1.13 | 0.3275 to 1.932 | Yes | ** |
| 7 dpi vs. 42 dpi | 0.798 | -0.004243 to 1.600 | No | ns |
| 12 dpi vs. 19 dpi | 0.3506 | -0.4517 to 1.153 | No | ns |
| 12 dpi vs. 42 dpi | 0.01888 | -0.7834 to 0.8212 | No | ns |
| 19 dpi vs. 42 dpi | -0.3317 | -1.134 to 0.4706 | No | ns |

**Table S3: Mixed design two way ANOVA fixed effects and post hoc results for male longitudinal data.**

| **Table S4** | **3 dpi** |
| --- | --- |

| Source of Variation | % of total variation | P value | P value summary | F (DFn, DFd) |
| --- | --- | --- | --- | --- |
| Interaction | 15.41 | 0.0242 | * | F (4, 23) = 3.437 |
| gene | 36.26 | 0.0003 | *** | F (4, 23) = 8.086 |
| sex | 19.97 | 0.0003 | *** | F (1, 23) = 17.82 |
| Holm-Šídák's multiple comparisons test | Predicted (LS) mean diff. | Below threshold? | Summary | Adjusted P Value |
| **Female- males** |  |  |  |  |
| cd11b | 5.868 | No | ns | 0.1322 |
| cd3 | 11.54 | Yes | ** | 0.0038 |
| cd19 | 9.659 | Yes | ** | 0.0083 |
| vegfa | -0.04738 | No | ns | 0.9998 |
| pdgfa | -0.00172 | No | ns | 0.9998 |

| Table S4 | **7 dpi** |
| --- | --- |

| Source of Variation | % of total variation | P value | P value summary | F (DFn, DFd) |
| --- | --- | --- | --- | --- |
| Interaction | 30.97 | 0.0031 | ** | F (4, 24) = 5.367 |
| gene | 37.6 | 0.0011 | ** | F (4, 24) = 6.517 |
| sex | 4.369 | 0.0946 | ns | F (1, 24) = 3.029 |
| Holm-Šídák's multiple comparisons test | Predicted (LS) mean diff. | Below threshold? | Summary | Adjusted P Value |
| **Female- males** |  |  |  |  |
| cd11b | -13.4 | Yes | *** | 0.0002 |
| cd3 | 0.3092 | No | ns | 0.9919 |
| cd19 | 0.9503 | No | ns | 0.9811 |
| vegfa | 0.01485 | No | ns | 0.9957 |
| pdgfa | 1.419 | No | ns | 0.9811 |

| Table S4 | **12 dpi** |
| --- | --- |

| Source of Variation | % of total variation | P value | P value summary | F (DFn, DFd) |
| --- | --- | --- | --- | --- |
| Interaction | 4.091 | 0.6607 | ns | F (4, 20) = 0.6092 |
| gene | 57.19 | 0.0003 | *** | F (4, 20) = 8.518 |
| sex | 5.144 | 0.0954 | ns | F (1, 20) = 3.064 |
| Holm-Šídák's multiple comparisons test | Predicted (LS) mean diff. | Below threshold? | Summary | Adjusted P Value |
| **Female- males** |  |  |  |  |
| cd11b | 4.631 | No | ns | 0.276 |
| cd3 | 1.37 | No | ns | 0.9183 |
| cd19 | 0.4525 | No | ns | 0.9772 |
| vegfa | 0.1329 | No | ns | 0.9772 |
| pdgfa | 2.604 | No | ns | 0.7322 |

| Table S4 | **19 dpi** |
| --- | --- |

| Source of Variation | % of total variation | P value | P value summary | F (DFn, DFd) |
| --- | --- | --- | --- | --- |
| Interaction | 29.22 | 0.0003 | *** | F (4, 20) = 8.797 |
| gene | 37.39 | <0.0001 | **** | F (4, 20) = 11.26 |
| sex | 16.79 | 0.0002 | *** | F (1, 20) = 20.22 |
| Holm-Šídák's multiple comparisons test | Predicted (LS) mean diff. | Below threshold? | Summary | Adjusted P Value |
| **Female- males** |  |  |  |  |
| cd11b | -0.1312 | No | ns | 0.9982 |
| cd3 | 24.23 | Yes | **** | <0.0001 |
| cd19 | -0.1854 | No | ns | 0.9982 |
| vegfa | 5.231 | No | ns | 0.4516 |
| pdgfa | 5.021 | No | ns | 0.4516 |

| Table S4 | **42 dpi** |
| --- | --- |

| Source of Variation | % of total variation | P value | P value summary | F (DFn, DFd) |
| --- | --- | --- | --- | --- |
| Interaction | 15 | 0.3225 | ns | F (4, 20) = 1.249 |
| gene | 21.35 | 0.1729 | ns | F (4, 20) = 1.778 |
| sex | 3.623 | 0.285 | ns | F (1, 20) = 1.207 |
| Holm-Šídák's multiple comparisons test | Predicted (LS) mean diff. | Below threshold? | Summary | Adjusted P Value |
| **Female- males** |  |  |  |  |
| cd11b | -2.393 | No | ns | 0.542 |
| cd3 | -2.363 | No | ns | 0.542 |
| cd19 | -0.9076 | No | ns | 0.8161 |
| vegfa | 1.809 | No | ns | 0.6024 |
| pdgfa | -0.01644 | No | ns | 0.9918 |

**Table S4: Two way ANOVA fixed effects and post hoc results for female and male comparative data.**
